# Molecular basis for anti-jumbo phage immunity by AVAST Type 5

**DOI:** 10.1101/2025.07.08.663546

**Authors:** Aswin Muralidharan, Ana Rita Costa, Desi Fierlier, Daan Frits van den Berg, Halewijn van den Bossche, Adja Damba Zoumaro-Djayoon, Martin Pabst, Martin Pacesa, Bruno E. Correia, Stan J. J. Brouns

**Affiliations:** Department of Bionanoscience, Delft University of Technology, van der Maasweg 9, 2629 HZ Delft, The Netherlands; Kavli Institute of Nanoscience, Delft University of Technology, van der Maasweg 9, 2629 HZ Delft, The Netherlands; Department of Biotechnology, Delft University of Technology, 2629 HZ Delft, Netherlands; Laboratory of Protein Design and Immunoengineering, École Polytechnique Fédérale de Lausanne and Swiss Institute of Bioinformatics, Lausanne, Switzerland

**Keywords:** Defense systems, *Pseudomonas aeruginosa*, Jumbo phage, Anti-jumbo phage immunity, AVAST, Sir2, AAA+ ATPase, STAND, TurboID, EPI vesicle

## Abstract

Jumbo phages protect their genomes from DNA-sensing bacterial defense systems by enclosing them within vesicles and nucleus-like compartments. Very little is known about defense systems specialized to counter these phages. Here, we show that AVAST Type 5 (Avs5) systems, part of the STAND superfamily and spanning three phylogenetic Avs5 clades, confer conserved immunity against jumbo phages. Using localization microscopy and biotin proximity labeling we demonstrate that Avs5 localizes to early infection vesicles, where it senses an essential, early expressed phage protein named JADA—Jumbophage AVAST5 Defense Activator. Recognition of phage infection triggers the Sir2-like effector domain of Avs5 across all three Avs5 clades, resulting in rapid NAD^+^ hydrolysis, disruption of phage nucleus formation, and arrest of infection. These findings reveal a spatially coordinated bacterial immune strategy that targets an early vulnerability in jumbo phage infection.

## Introduction

Bacteriophages of the *Timaliidae* family (also known as nucleus forming jumbo phages) have evolved to possess an innate capability to elude detection by the prokaryotic immune systems like DNA targeting CRISPR-Cas systems and restriction modification systems^1–3^. To do so, they follow a carefully coordinated infection life cycle and sequential compartmentalization. During the early stages of infection, these phages compartmentalize their genome using an early phage infection (EPI) vesicle and evade detection by DNA sensing defense systems^4–7^. This evasion is achieved in middle and late stages of infection through the physical concealment of their genome from bacterial cytoplasm within a selectively permeable proteinaceous nucleus-like genome replication and transcription compartment^8–11^. This nucleus is enclosed by a self-assembling crystalline lattice (nuclear shell) composed primarily of the phage encoded protein ChmA and is centered inside the bacterium with the help of phage encoded tubulin like filament PhuZ^10,12,13^. The phage injects a multi-subunit virion RNA polymerase (vRNAP) packaged in the virion along with the genome into the EPI vesicle to transcribe the early viral genes, and a DNA polymerase, presumably for replication^14–16^. To transcribe the late viral genes once in the nuclear shell, *Chimalliviridae* also carry a multi-subunit non-virion RNA polymerase (nvRNAP) composed of early gene products^14,17,18^. The transcription of these phages is hence independent of the host RNA polymerases^19^.The phage mRNA is transcribed inside the nucleus and is then translated outside the nucleus in the bacterial cytoplasm, implying it is trafficked through the nuclear shell^8,9,11,20,21^. The phage procapsids assemble outside the nucleus, and then treadmill over PhuZ to dock onto the nuclear shell where the genome is packaged into the capsid^22,23^. The mature capsids then detach from the nuclear shell and accumulate with the assembled phage tail components to form mature phage particles^8^. The host bacterium is then lysed, and the phage particles are released to infect new cells. Since the phage genome is concealed by the nucleus from the start of the infection till the end, the host must rely on defense systems which detect either phage mRNA or proteins to activate an immune response.

The strategies bacteria use to combat *Chimalliviridae* involve initiating immune responses through their adaptive and innate immune systems. While the phage DNA is protected in the nuclear shell from DNA targeting CRISPR-Cas systems (Type I, II and VI), the RNA targeting CRISPR-Cas systems (Type III or Type VI) are effective by recognizing specific mRNA nucleotide sequences that correspond to phage transcripts in the bacterial cytoplasm^1,2,24,25^. This strategy involves activation of effectors that lead to host growth arrest or cell death, such that phage propagation is inhibited^24^. Only a few innate defenses effective against *Chimalliviridae* have been identified^26^, including Juk, the only characterized defense system specific to jumbo phages^27^.

AVAST Type 5 or Avs5 belongs to the ancient signal transduction ATPases of numerous domains (STAND) NTPase superfamily which play a crucial role in pathogen associated molecular pattern (PAMP) triggered immunity across all domains of life^28–36^. The STAND NTPases share a common tripartite modular domain architecture, featuring a central NTPase domain, a C-terminal sensor, and an N-terminal effector. The variable effector domain is responsible for initiating programmed cell death when the sensor recognizes PAMPs. STAND proteins in plants and animals are capable of dynamically transitioning between on and off states to regulate activity upon recognition of PAMPs, cytokines and damage associated molecular patterns^37–41^. With evolutionary and structural similarities to universally used nucleotide-binding oligomerization domain (NOD)-like receptors (NLRs), Avs proteins represent a common link in the pattern recognition immune mechanisms between bacteria and other domains of life^31,32,35,42–45^. The nuclease activity of Avs1-Avs4 lead to altruistic cell death by cleaving both the phage and host genome. This occurs by recognizing conserved structural patterns within the phage DNA packaging machinery, namely the large subunit of phage terminase and portal proteins or another phage protein Ksap1 of unknown function^29,35^. By direct recognition of these highly conserved and essential phage proteins, all characterized Avs systems provide the bacterial host immunity against a broad range of phages^29,33,36,46^. In contrast to the nuclease-based effectors of Avs1–4, Avs5 systems employ a Sir2 NADase. It remains an open question what activates Avs5 systems and what governs their phage specificity. Hence, we systematically studied the phylogeny, activation and molecular mechanisms of Avs5 immunity.

We show that Avs5 systems form three distinct phylogenetic clades, each conferring conserved immunity against nucleus-forming jumbo phage Pa36. One clade additionally mediates broad-spectrum defense, targeting phages ranging from ssRNA viruses to members of the *Mesyanzhinovviridae*. We show that jumbo phage-specific *Pseudomonas aeruginosa* Avs5 (PaAvs5-1) system localizes to the EPI vesicle. Using proximity labelling, we identified that PaAvs5 from all the three clades recognize an early-expressed, essential *Chimalliviridae*-specific protein, Pa36 gp316—here named JADA (Jumbo phage Avs5 Defense Activator). Cryo-EM analysis reveals that JADA forms a homodimer. Upon infection with a jumbo phage, these recognition events initiate NAD⁺ hydrolysis by the PaAvs5 Sir2 domain, disrupting a key metabolic cofactor. We further show that using this metabolic disruption, PaAvs5 disrupts jumbo phage nucleus formation and slows progeny production. Together, our findings uncover the molecular basis, spatial localization, and temporal activation of AVAST Type 5 antiviral defense systems against jumbo phages.

## Results

### Shared Anti-Jumbo Phage Immunity by Avs5 Homologs Across Phylogenetic Clades

Recent studies showed that Avs5 systems, despite sharing a conserved domain architecture (Sir2-STAND) exhibit striking diversity in their anti-phage specificity across bacterial species^30,36^. For example, the Avs5 system from *E. fergusonii* (EfAvs5) confers broad protection against diverse phages in its expression host^36^, whereas an *E. coli* Avs5 homolog (EcAvs5) provides targeted immunity against the T2 phage^30^. Similarly, *P. aeruginosa* Avs5 (PaAvs5) specifically defends against the nucleus-forming jumbo phage Pa36^26^. These contrasting immunity profiles, despite shared domain architecture, raise a key question: why do closely related Avs5 systems exhibit such divergent anti-phage specificities?

To understand this contrasting anti-phage immunity, we performed a phylogenetic classification of Avs5 homologs with a Sir2 effector domain. An unrooted phylogenetic tree revealed three distinct clades: PaAvs5 is found in Clade 1 (PaAvs5-1), EcAvs5 in Clade 2, and EfAvs5 in Clade 3 (Figure 1A). To determine the range of phages against which Avs5 homologs confer immunity, we selected one representative homolog from each clade in our clinical *P. aeruginosa* collection (Figure 1A,B). These homologs were cloned into *P. aeruginosa* PAO1 using the low copy pUCP20 plasmid under their native promoters. We then exposed these strains to a diverse panel of 18 phages from 7 families, including those from *Autographiviridae, Mesyanzhinovviridae*, *Bruynoghevirus*, *Casadabanvirus, Pbunavirus*, an ssRNA *Fiersviridae* phage, and one *Chimalliviridae* jumbo phage, all capable of infecting PAO1^26^. Avs5 homologs from Clades 2 (PaAvs5-2) and 3 (PaAvs5-3) provided immunity against the ssRNA phage PP7, while PaAvs5-3 additionally protected against *Mesyanzhinovviridae* phage Pa53 (Figure 1F). Notably, all tested Avs5 systems conferred strong immunity against the nucleus-forming jumbo phages Pa36, reducing its efficiency of plating by over five orders of magnitude (Figure 1E). PaAvs5-1 displayed exclusive immunity to jumbo phage.

**Figure 1.**
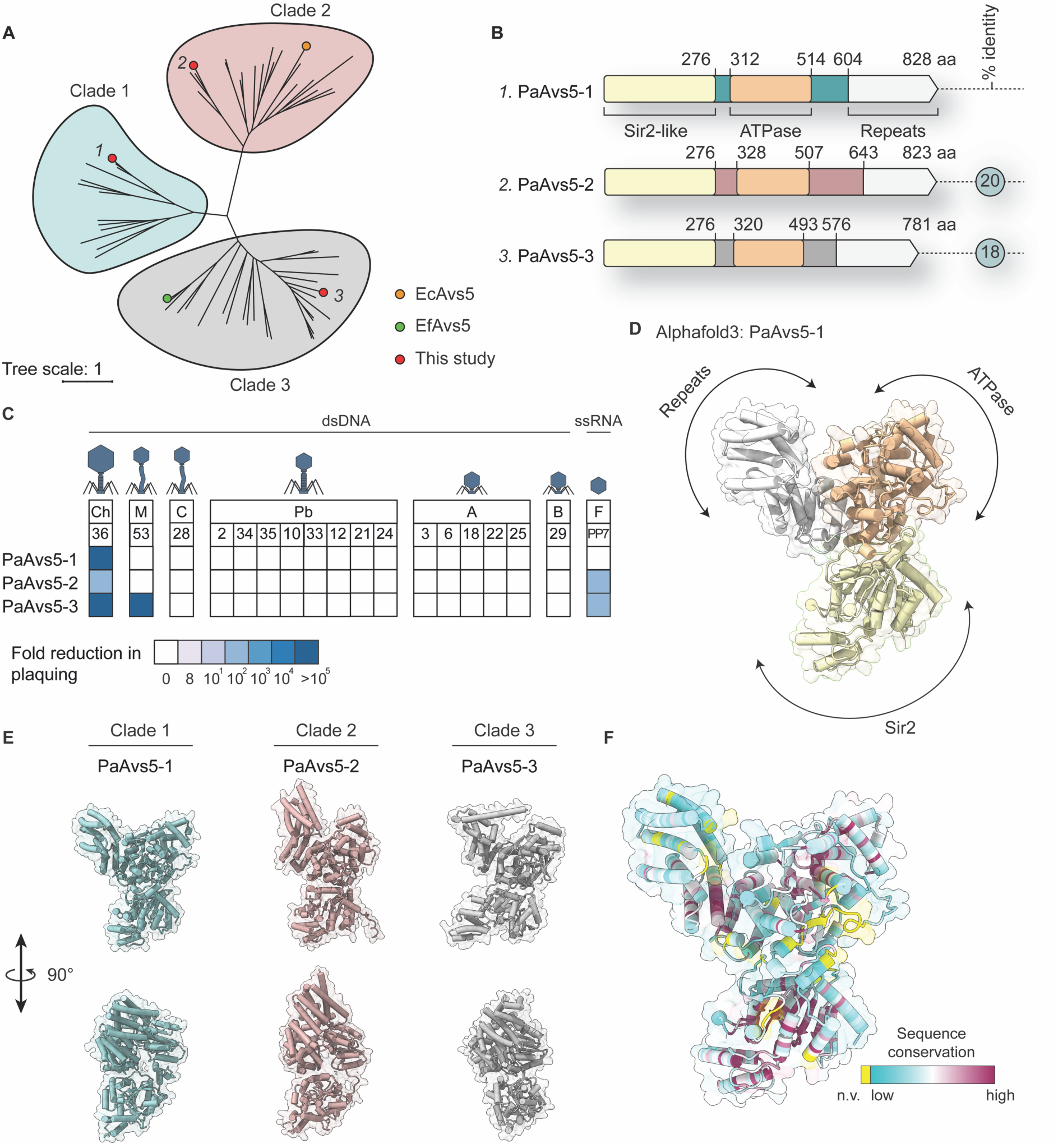
Phylogenetic distribution, domain architecture, and anti-phage immunity of Avs5 systems. (A) Phylogenetic tree of Sir2-STAND Avs5 homologs, grouped into three major clades. EcAvs5 (orange) and EfAvs5 (green) are included as references. The three PaAvs5 variants analyzed in this study are highlighted in red. The selected homologs are named with the clade number in superscript (i.e., PaAvs5-x is from clade x) (B) Domain architectures of the three tested PaAvs5 systems, with domain boundaries indicated. Superscripts correspond to the clade assignment in (A). (C) Phage immunity profiles of PaAvs5 systems, shown as fold reduction in plaquing across a panel of phages from different families and genera: A: (*Autographiviridae*), B (*Bruynoghevirus*), M (*Mesyanzhinovviridae*), F (*Fiersviridae*), Pb (*Pbunavirus*), Ch (*Chimalliviridae*), C (*Casadabanvirus*). (D) AlphaFold3 predicted structural model of PaAvs5-1, highlighting the Sir2 (in yellow), ATPase (in orange), and repeat-containing domains (in white). (E) Alphafold3 predicted structural comparison of PaAvs5 homologs from each clade (PaAvs-1, PaAvs5-2, PaAvs5-3). (F) Sequence conservation mapped onto the PaAvs5-1 Alphafold3 structure, based on a multiple sequence alignment of 64 PaAvs5 homologs. Coloring follows a cyan-to-maroon gradient, with highly conserved residues in maroon and variable regions in cyan. Residues without a conservation value (n.v.), including unaligned positions, are shown in yellow.

Hence, our findings reveal striking plasticity in the anti-phage immunity profiles of Avs5 systems. In particular, the PaAvs5-3 conferred immunity against a remarkably diverse phage genera, including the ssRNA phage PP7, which carries only four coding sequences within a ∼3.6 kb genome, a siphophage with an ∼62 kb genome encoding 89 coding sequences, and a jumbo phage with a 280 kb genome encoding over 300 coding sequences. This variation mirrors previous observations on Avs5 systems in *E. coli* and *E. fergusonii*, where representative homologs conferred targeted or broad-spectrum immunity, respectively^30,36^. While Avs5 homologs from different phylogenetic clades vary in the breadth of their phage targets, all retain conserved activity against nucleus-forming jumbo phages revealing a shared core specificity.

### Structural and Sequence Variability in Avs5 Sensing Domains Potentially Drive Immunity Differences

To understand the molecular basis of Avs5 immune response, we checked its domain architecture (Figure 1B). Consistent with other Avs defense systems, PaAvs5 we investigated shared a tripartite domain architecture. At the amino-terminus, they are predicted to contain a silent information regulator 2 (Sir2)-like effector domain (Pfam: PF13289), nested within a DHS-like NAD/FAD-binding domain superfamily (SSF: SSF52467)^47,48^. Additionally, they possess a novel STAND NTPase3 subdomain (nSTAND3, Pfam: PF20720) situated within an AAA+ ATPase domain (SMART: SM00382)^28,49,50^. The carboxy terminal repeat domains in the tested Avs5 systems are predicted to contain different types of structural repeats, including tetratricopeptide repeats (TPR) and Sel1-like repeats in PaAvs5-1 and PaAvs5-2; and pentatricopeptide repeats (PPR) in PaAvs5-3. These superstructural alpha-helical repeat-based domains are often implicated in protein ligand binding^51,62^.

Pairwise alignments between tested homologs indicate low sequence identity across Avs5 homologues, with PaAvs5-1 and PaAvs5-2 sharing 20% identity (35% similarity), PaAvs5-1 and PaAvs5-3 sharing 18% identity (35% similarity), and PaAvs5-2 and PaAvs5-3 sharing 21% identity (35% similarity). Analysis of amino acid sequence conservation among homologs from our clinical *P. aeruginosa* collection revealed poor conservation in the region between the Sir2 and nSTAND3 domains, the ATPase domain, and the carboxyl-terminal domain consisting of repeating alpha helices (Figure 1F, S1A, S1B).

Alphafold3 structural predictions suggest a potentially “closed” state, where the three domains adopt a compact configuration stabilized by extensive intradomain interactions. (Figure 1C). In this predicted structure, the repeat domain appears to fold back on to the ATPase domain, possibly to block the oligomerization mediated by this domain. Structural predictions using AlphaFold3 and subsequent structural alignments revealed subtle variations among tested homologs, particularly in the organization of the sensing domain (Figure 1C, 1D). Pairwise structural alignments show similar folds, with PaAvs5-2 and PaAvs5-3 displaying TM-align scores of 0.68 and 0.67, respectively, in comparison to PaAvs5-1.

Outside prokaryotes, the predicted structure of PaAvs5 shows strong similarity to Sterile alpha motif domain-containing protein 9/SAMD9 (Uniprot: Q5K651) from *Homo sapiens*. (Foldseek ^52^ BFMD database: probability = 1, E-value = 4.6 x 10^-16^, position in query = 3-818). Additionally, there is strong structural homology to NB-ARC domain-containing protein from *Cladophialophora carrionii* (Uniprot: A0A1C1CV94, Foldseek: probability = 1, E-value = 3.7 x 10^-6^, position in query = 199-825). These proteins are implicated in innate immune responses, including antiviral roles^53,54^. Despite low sequence identities (<15%), these structural homologies suggest a shared cross-kingdom functional similarities.

Our results suggest that variability in both the sequence and structure of the carboxy-terminal repeat regions and the region between the Sir2 and nSTAND3 domains of the Avs5 homologs could play a role in recognizing distinct phage ligands. These structural differences may explain the diverse immunity profiles observed across the Avs5 clades. Given PaAvs5-1’s exclusive protection against the jumbo phage Pa36, we next sought to elucidate the molecular basis of its activation and immune response.

### PaAvs5 Restricts Phage Propagation and Delays Jumbo Phage Replication

To determine how PaAvs5-1 protects against Pa36, we infected strains with Pa36 at different multiplicities of infection (MOI) in liquid culture. In the absence of infection, both control and PaAvs5-1 strains grew similarly (Figure 2A). At a MOI = 0.01, the PAO1 strain carrying PaAvs5-1 grew unaffected, while the control strain with an empty plasmid experienced culture collapse within four hours (Figure 2B). However, at a MOI = 1, both strains underwent culture collapse (Figure 2C). Additionally, PaAvs5-1 strains reduced Pa36 plaquing efficiency by seven orders of magnitude compared to control strains (Figure 2D). Together, these results suggest that PaAvs5-1 may limit phage replication, possibly through altruistic cell death to restrict infection.

**Figure 2.**
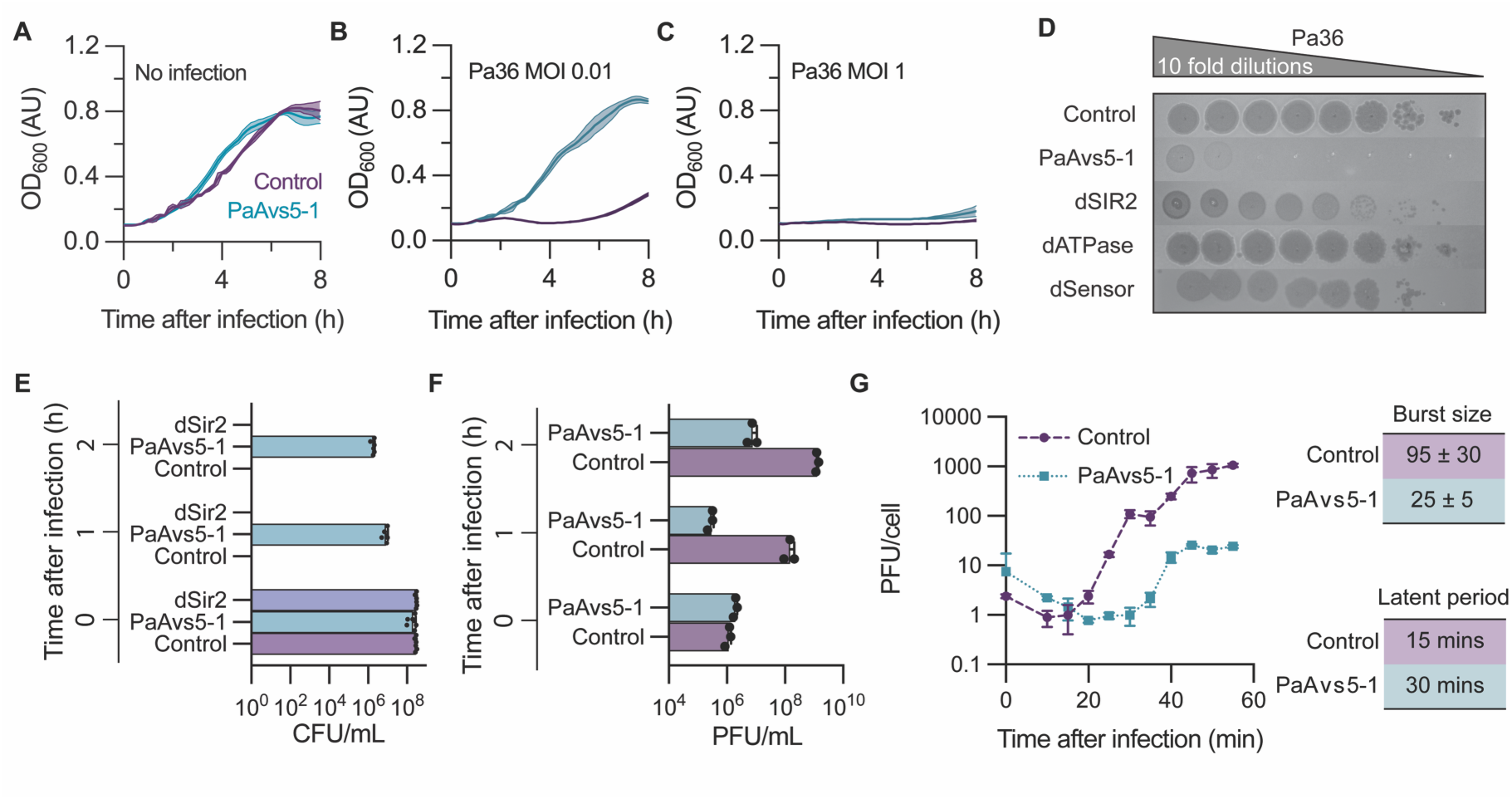
Jumbo phage resistance, and infection kinetics of PaAvs5. (A–C) Growth curves of PAO1 strains expressing PaAvs5-1 or a control (empty plasmid), in liquid culture with or without Pa36 infection. (A) No infection control showing similar growth. (B) Infection with Pa36 at MOI = 0.01, demonstrating immunity provided by PaAvs5-1, while the control culture collapses. (C) Infection with Pa36 at MOI = 1, resulting in culture collapse in both PaAvs5-1-expressing and control cells. (D) Efficiency of plaquing assay for Pa36 on wild-type PaAvs5-1 and inactivating mutants. PaAvs5-1 cells show seven-log protection against Pa36 compared to control cells. dSir2 (N110A), dATPase (K363A), and dSensor (Δ812–828) are unable to provide protection. (E) Bacterial survival measured in colony forming units/mL (CFU/mL) at different time points after infection by Pa36 (MOI = 10). (F) Phage replication measured in plaque forming units/mL (PFU/mL) at different time points after infection by Pa36 (MOI = 0.1). (G) One-step growth curve of Pa36 on control and PaAvs5-1-expressing strains, showing burst size and latent period.

To assess whether PaAvs5-1 protects the host bacterium, we measured colony-forming units (CFU/mL) after infecting PAO1 with Pa36 at a MOI = 10 for different durations (Figure 2E). PAO1 strains lacking functional PaAvs5-1 had no surviving colonies after infection for one hour, indicating complete lysis. PAO1 strains carrying PaAvs5-1 showed only a ∼2-log reduction in CFU even after infection for two hours, demonstrating that PaAvs5-1 provides protection under these conditions.

To assess whether PaAvs5-1 activity affects phage replication, we measured plaque-forming units (PFU) after infecting the strains at MOI = 0.1 (Figure 2F). In control cells, phage replication increased from 10^6^ PFU/mL before infection to nearly 10^9^ PFU/mL within two hours. In contrast, PaAvs5-1-containing cells exhibited a reduced increase in phage titers during the first hour. After two hours, the titers in these cells showed only a modest increase, reaching 10^7^ PFU/mL from an initial 2 × 10^6^ PFU/mL.

Next, we performed a one-step growth curve analysis to test the single replication cycle of the phage (Figure 2G). In the absence of PaAvs5-1, phage release (latent period) began approximately 15 minutes post-infection. However, in the presence of PaAvs5-1, the latent period doubled. The burst size or phage particles released per infected cell was reduced fourfold in PaAvs5-1 strains compared to controls. Combined, these effects resulted in an almost 100-fold reduction in total phage release one-hour post-infection (Figure 2F, 2G).

Altogether, we demonstrate that PaAvs5-1 provides the host bacterium immunity by slowing jumbo phage propagation and reducing the efficiency of phage replication.

### Sir2 Domain of PaAvs5 Initiates NAD^+^ Hydrolysis Upon Activation

The Sir2-like domain in PaAvs5-1 shares sequence similarity with the Sir2-like domains of ThsA from the Thoeris defense system and the DSR2 defense system, both of which hydrolyze NAD^+^ to trigger antiphage activity^55,56^. The proposed catalytic mechanism, like other Sirtuins, involves Asn110 coordinating a water molecule to position it near the ribose ring oxygen adjacent to the nicotinamide (Nam) ring of NAD^+^ (Figure 3A)^57^. Subsequently, this is expected to facilitate the hydrolysis of NAD^+^ to adenosine 5′-diphosphoribose (ADPR) and Nam upon activation.

**Figure 3.**
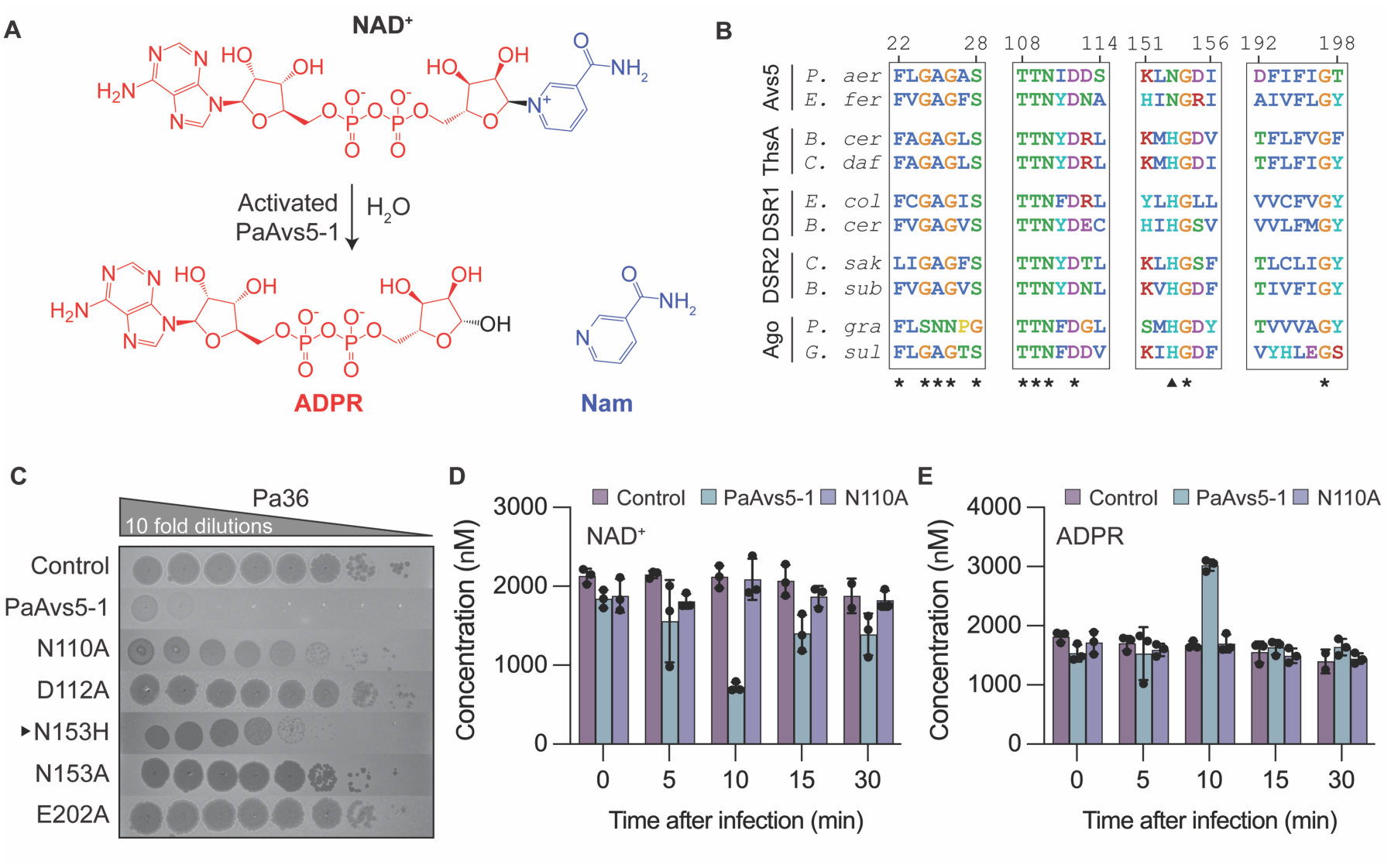
The Sir2 domain of PaAvs5 hydrolyses NAD^+^ upon activation. (A) Schematic of NAD^+^ hydrolysis by activated PaAvs5-1, generating ADPR and Nam. (B) Multiple sequence alignment of the Sir2 domain across Avs5, ThsA, Ago and DSR1/DSR2 homologues from diverse bacterial species. Conserved residues are marked (*). Amino acids are colored based on CLUSTAL color scheme. (C) Efficiency of plaquing assay showing the impact of Sir2 domain mutations on PaAvs5-1-mediated defense against Pa36. Tenfold dilutions of Pa36 were spotted on PAO1 strains expressing wild-type, or mutant PaAvs5-1, or empty plasmid control. (D, E) Intracellular NAD^+^ (D) and ADPR (E) concentrations measured in PAO1 cells infected with Pa36 (MOI = 3) at indicated time points post-infection. Infection was initiated when cultures reached OD600 = 0.3. Data represent mean ± SD (n = 3 biological triplicates), with individual data points shown.

Amino acid sequence alignments of the Sir2-like domains from Avs5, DSR1, DSR2, ThsA, and short prokaryotic Argonautes (Ago) reveal the presence of highly conserved motifs, GAGXS (a.a. 24–28 in PaAvs5-1) and TTNXD (a.a. 108–112), which are characteristic of all known Sirtuins (Figure 3B, S1B). These motifs were previously shown to directly interact with NAD^+^ in Sirtuins^58^. Site directed point substitutions in the predicted catalytic residues to alanine in the predicted Sir2-like domain (N110A, D112A, E202A) abrogated the immunity imparted by PaAvs5-1 (Figure 3C, S2). This shows that the catalytic NAD^+^ hydrolysis activity of the Sir2 domain is essential for the PaAvs5-1 immune response.

Surprisingly, PaAvs5-1 lacks the canonical catalytic HG motif, and instead harbors an NG (a.a. 153-154) motif in its place. The HG motif, with the catalytic histidine is considered to be absolutely conserved in Sir2 domains across domains of life^58–60^. We found that the NG motif occurs nearly as frequently as the canonical HG motif among Avs5 homologs, appearing in approximately a one-to-one ratio (Figure S1C). In eukaryotic Sirtuins, this motif couples NAD^+^ hydrolysis to ADP-ribosylation or deacetylation of protein substrates containing acetylated lysine residues^57^. Substituting Asn153 in PaAvs5-1 with histidine partially maintained activity but significantly reduced immunity (Figure 3C). Alanine or glutamine substitution of Asn153 abolished protection, indicating that Asn153 or His153 is required for defense, with Asn153 being more efficient (Figure 3C). The NG motif therefore is a novel variant of the canonical HG motif in the Sir2 domain.

To investigate the phage-activated NAD^+^ hydrolysis activity of the Sir2 domain in PaAvs5, we measured NAD^+^ and ADPR concentrations in Pa36 infected cells (MOI = 3, OD_600_ = 0.3) at various time points post-infection. In PAO1 cells either lacking PaAvs5-1 or expressing a PaAvs5-1 variant with an inactive Sir2-like domain (N110A), NAD^+^ (∼2100 nM) and ADPR (∼1800 nM) levels remained stable during the first 30 minutes following Pa36 infection (Figure 3D, 3E). In contrast, cells expressing functional PaAvs5-1 exhibited a threefold decrease in NAD^+^ (∼700 nM) 10 minutes post-infection, accompanied by over 50% increase in ADPR (∼3000 nM). This demonstrates the successful hydrolysis of the nicotinamide-ADPR bond of NAD^+^ by the Sir2 domain to produce ADPR.

However, to our surprise, NAD^+^ concentrations in PaAvs5-1 expressing cells rebounded to basal levels within 15 minutes post-infection and remained stable through 30 minutes post-infection (Figure 3D). Similarly, ADPR concentrations returned to basal levels (∼1600 nM) during the same period (Figure 3E). This suggests that ADPR may be recycled back into NAD^+^ by either jumbo phage-encoded NAD^+^- recycling proteins or host enzymes, but the exact cause is currently unknown. Nonetheless, the temporary reduction of the NAD^+^ pool appears sufficient for PaAvs5-1 to hinder Pa36 from successfully infecting the cell. Overall, our data suggests that the catalytic activity of Sir2 domain is essential for the antiphage defense of PaAvs5.

### ATPase Activity of PaAvs5 is Essential for Immunity

The AAA+ ATPase domain in PaAvs5-1 contains a phosphate-loop (P-loop) NTPase subdomain that often adopts a Rossmann fold and includes a Walker A motif (GPAGSGKT, amino acids 357–364, consensus Walker A motif: GX_4_GK[S/T] with X representing any amino acids)^49,61,62^. This glycine rich motif with an invariant lysine is commonly involved in the phosphate binding of ATP or GTP, with the Gly362, Lys363, and Thr364 residues in PaAvs5-1 coordinating the position of the triphosphate group (Figure S1B)^28,49^. Replacing the Gly362 or Lys363 to an alanine abrogated the PaAvs5-1 immunity (Figure 2D, S2). This indicates that ATP phosphate binding by the Walker A motif is essential for PaAvs5-1-mediated immunity.

Secondary structure predictions and AlphaFold3 modelling suggest that PaAvs5-1 contains a putative Walker B-like motif (IIFIE, residues 406–410). Walker B motifs are conserved elements in P-loop NTPases, typically consisting of four hydrophobic residues followed by two consecutive acidic residues^61^. In PaAvs5-1, this motif has a single acidic residue (Glu410) instead of the canonical h_4_D[D/E] Walker B motif, where h represents any hydrophobic residue (Figure S1D). In PaAvs5-1, Arg411 is present within the ATPase pocket in place of a second acidic residue (Figure S1D). These residues are known to coordinate a catalytically essential Mg^2+^ ion for ATP hydrolysis. Mutating Glu410 to Ala abolishes the protective phenotype while mutation of Glu410 to Gln retained protective phenotype (Figure S1F). While the carboxylate group in Glu can coordinate Mg^2+^ directly and participate in proton abstraction to perform ATP hydrolysis, amide group of Gln lacks these capabilities. Together, these findings suggest that the Walker B motif in PaAvs5-1 primarily facilitates ATP binding and conformational changes rather than directly catalyzing ATP hydrolysis. Additionally, Arg438, a highly conserved residue at the carboxy-terminal tip of β12, is also essential for immunity. Its substitution to Ala eliminates protection against Pa36 (Figure S1F). Arg438 potentially serves a role as an Arginine finger, often involved in forming contacts with γ-phosphate of nucleotide.

These findings indicate that nucleoside triphosphate binding or hydrolysis by these residues is necessary for PaAvs5-1. Overall, our results show that ATPase domain activity is critical for protection. Specifically, the Walker A and Walker B motifs, along with the Arginine finger—key elements involved in nucleotide binding—are required for PaAvs5 function.

### C-Terminal Repeat Domain of PaAvs5 is Essential for Activation

To understand how PaAvs5 senses the PAMP, we examined its carboxy-terminal repeat domain, which likely serves a dual function: maintaining PaAvs5 in an inactive state while also acting as a pathogen sensor. Within this region, we identified two predicted repeat motifs in PaAvs5-1: one TPR motif (672–705) and one Sel1 motif (residues 713–748)^63,64^. AlphaFold3 structural predictions indicate that each motif adopts a tandem helix-turn-helix structure, with helices arranged in an antiparallel fashion. Each helix within these repeats, typically 13 to 20 amino acids in length. Repeat domains, such as these, are commonly involved in protein-protein interactions^51,65^. Despite being poorly conserved, deletion of any predicted helices in the C-terminal region (PaAvs5-1 Δ812–828, PaAvs5-1 Δ795–828, or PaAvs5-1 Δ769– 828) completely abolished PaAvs5-1-mediated immunity towards Pa36 (Figure 1D, Figure S1F). Hence, this variable region near the C-terminus plays a critical role in sensing phage infection and activating the defense response.

Towards the amino terminus of the repeat domain, we identified two highly conserved aromatic residues—Tyr661, Trp675, and an invariant Gln677 (Figure S1B, S1E). Within the AlphaFold3 structural predictions, these residues cluster on consecutive anti-parallel α-helices within the repeat domain (Figure 1F). Its strict conservation and location bridging the TPR and ATPase domains suggest a role in serving as a pivot point for conformational switching.

### PaAvs5 Inhibits Formation of Jumbo Phage Nucleus, and it Migrates to the Phage Infection Site

To study PaAvs5 localization and its effect on phage infection, we expressed a C-terminal mNeonGreen-tagged PaAvs5-1 from its native promoter in *P. aeruginosa* PAO1 using the pCUP20 plasmid. This fusion retained full protective activity against phage Pa36 (Figure S1F). Cells were infected at a high MOI =5, and both bacterial and phage DNA were visualized using DAPI staining. At 50 mpi, ∼60% of cells expressing the inactive PaAvs5-1 N110A mutant displayed distinct phage nuclei. In contrast, fewer than 5% of wild-type PaAvs5-1 cells formed a nucleus; instead, ∼80% showed localized DNA foci, typically at the poles, indicating an immune response early in infection. By 90 mpi, PaAvs5-1 N110A-expressing cells showed extensive DNA degradation and lysis, while PaAvs5-1-expressing cells maintained intact genomes and largely lacked nuclei (∼10% contained nuclei).

In line with current models of PhiKZ-like jumbo phage infection, the process initiates at the cell poles, where an early infection vesicle is formed in the host. Here, transcription of early phage genes occurs, whose transcripts are exported into the cytosol for translation. Our findings indicate that PaAvs5-1 detects these early polar-localized phage proteins, triggering its immune response. As a result, the infection is arrested at an early stage: in most cells, DNA foci remain localized at the poles, and phage genome trafficking in the nucleus is blocked. This suggests that key early processes involving PhuZ and ChmA, which are essential for transporting the phage genome to mid-cell and assembling the nucleus are effectively inhibited by PaAvs5-1. To determine where the PaAvs5-1 system localizes during infection, we examined its subcellular distribution. In uninfected cells, the PaAvs5-1-mNeonGreen signal was diffuse throughout the cytosol (Figure 4A, 4B, S2A-D). Upon phage infection, however, the signal was consistently excluded from the phage nucleus, regardless of whether the system was wild-type or catalytically inactive (Figure 4A, 4B, S2A-D). Strikingly, in ∼40% of infected cells (n = 300), PaAvs5-1 formed one or more bright foci by 50 minutes post-infection, often colocalizing with intense DNA foci—likely representing phage genomes. They were typically located at the cell poles and along the membrane. (Figure 4A, 4E, S2D). Of these, ∼27% had a single focus and ∼13% had two or more (Figure 4E). A similar pattern was observed in cells expressing the catalytically inactive dSIR2 mutant: ∼50% of infected cells (n = 131) showed bright foci, again localized to the poles and membrane, with ∼30% showing one focus and ∼20% showing multiple (Figure 4A, 4E, S2A). Interestingly, this polar and membrane-associated localization persisted even in sensor deletion (Δ769–828) and ATPase-inactive (K363A) mutants following Pa36 infection (Figure S2B, S2C), suggesting that recruitment of PaAvs5-1 to the site of infection occurs independently of its sensing and ATPase domains. Each focus is approximately three to four times brighter than the surrounding cytosol, suggesting the recruitment of PaAvs5-1 to the infection site (Figure S2E-H). In the inactive mutants, this intensity increases even further, potentially due to sustained production of phage-derived triggers that continue to be sensed (Figure S2E-H). These foci were absent during infection with Pa34, a phage that PaAvs5-1 cannot provide immunity against, implying that focus formation reflects a functional spatial organization required for antiviral defense (Figure S3).

**Figure 4.**
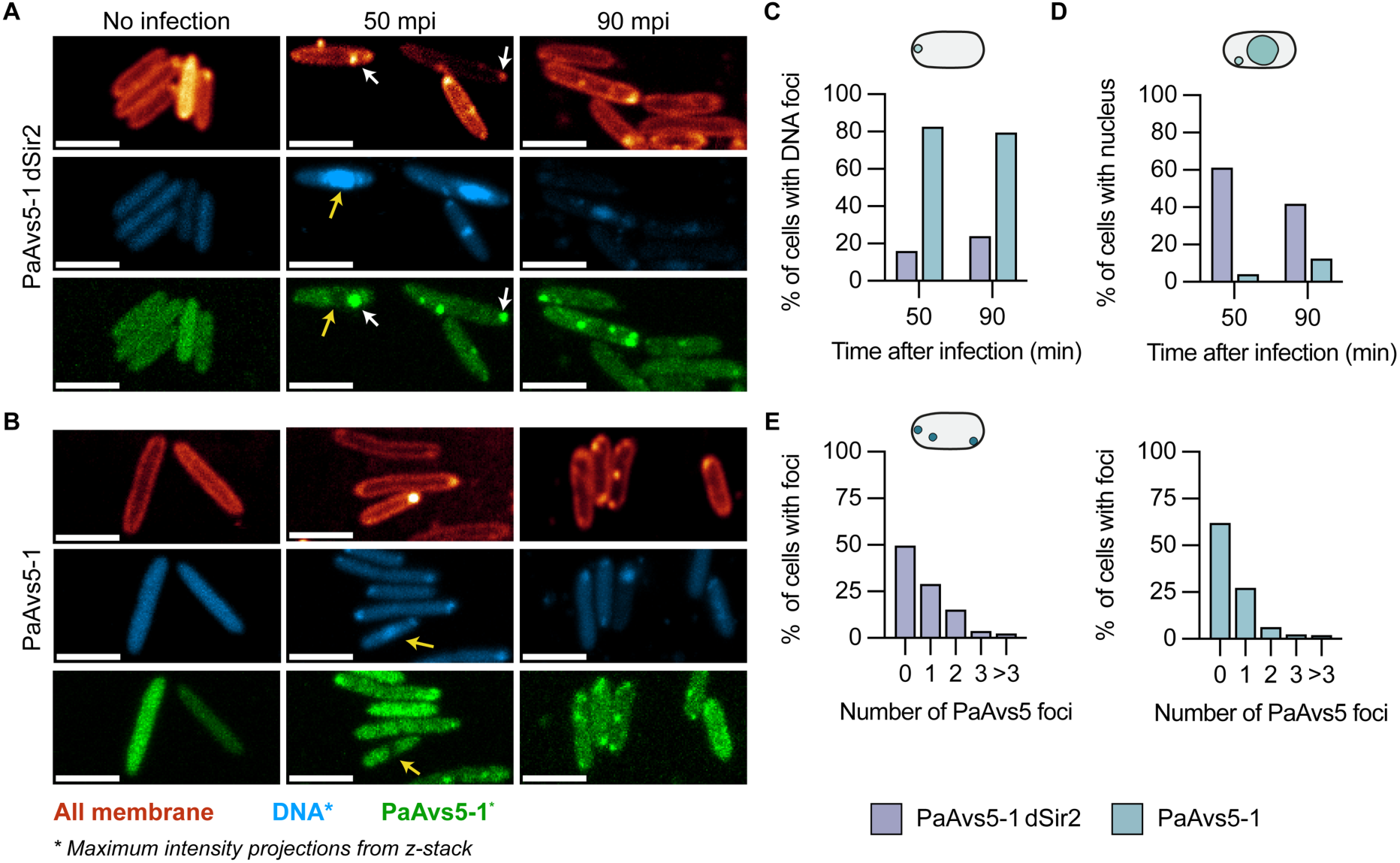
PaAvs5 inhibits formation of the jumbo phage nucleus and forms intracellular foci. (A, B) Confocal microscopy images of jumbo phage-infected *P. aeruginosa* PAO1 strains expressing mNeonGreen-tagged (green channel) (A) PaAvs5-1 N110A and (B) PaAvs5-1. The cell membrane is labelled with MitoTracker Deep Red-FM (orange channel), and DNA is stained with DAPI (blue channel). Representative PaAvs5-1 foci are indicated by white arrows. A yellow arrow highlights a phage nucleus and the corresponding region where PaAvs5-1 is excluded in the green channel. Phage infection proceeds to initiate cell lysis at 90 mpi in PaAvs5-1 dSir2. Each panel includes uninfected cells and cells at 50- and 90-minutes post-infection (mpi) shown as separate columns. DAPI and mNeonGreen-PaAvs5-1 channels are shown as maximum intensity projections from z-stacks; the membrane channel is shown as a single optical section to highlight clear cell outlines. Contrast in the DAPI channel of uninfected cells was adjusted to improve visibility. Scale bar: 3 μm. (C) Quantification of the percentage of infected cells showing visible DNA spots but no nucleus at 50 and 90 mpi for strains expressing either PaAvs5-1 dSir2/N110A (n = 131 cells at 50 mpi, 67 at 90 mpi) or wild-type PaAvs5-1 (n = 300 at 50 mpi, 137 at 90 mpi). Presence of DNA localization spots is estimated from the maximal intensity projections from z-stacks (as shown in schematic). (D) Quantification of the percentage of infected cells (from same cells in C) showing visible phage nuclei at 50 and 90 mpi for strains expressing PaAvs5-1 N110A or wild-type PaAvs5-1. Presence of nucleus is estimated from the maximal intensity projections from z-stacks (as shown in schematic). (E) Quantification of the percentage of infected cells (from the same dataset shown in C) displaying visible PaAvs5 foci at 50 minutes post-infection (mpi) in strains expressing either PaAvs5-1 N110A or wild-type PaAvs5-1. Foci were identified from maximum intensity projections of z-stacks, as illustrated in the schematic. Larger field of view microscopy images, along with additional data from dSensor and dATPase mutants, are provided in Figure S2A-D. Raw microscopy image data have been deposited at https://doi.org/10.5281/zenodo.15831563.

To assess whether these foci associate with early phage-induced (EPI) vesicles, we stained infected cells expressing PaAvs5-1-mNeonGreen with MitoTracker Deep Red-FM. This dye non-specifically labels cellular membranes, including internal membrane structures such as the EPI vesicle. Confocal imaging revealed frequent colocalization between PaAvs5-1 foci and MitoTracker signal, supporting the idea that PaAvs5-1 is recruited to membrane-associated compartments during early infection (Figure 4A, 4B). This association further strengthens our model of recruitment of PaAvs5-1 to the EPI vesicle, where early phage proteins are delivered and may be sensed by the defense system.

To investigate the localized distribution of PaAvs5-1 in infected cells we carefully inspected the amino acid sequence. The first 30 residues of its Sir2-like domain contain a predicted hydrophobic localization sequence, likely directing the protein to (intra)-cellular membranes. We hypothesized that this sequence facilitates PaAvs5-1’s interaction with early infection vesicles for proximity to the infection site^66,67^. Deleting the full localization sequence (Δ2–30) or just its N-region (Δ2–15) abolished protection (Figure S1F). Substituting a non-conserved residue in this helix (R9A) also disrupted protection. Hence, N-terminal localization sequence appears critical for effective phage sensing and defense.

In summary, PaAvs5-1 mounts a spatially organized immune response by localizing to membrane associated sites of phage entry. This spatial organization occurs independently of its catalytic or sensing functions. Upon infection, PaAvs5-1 accumulates at the poles and membrane, where it likely senses early phage proteins and blocks genome trafficking, preventing nucleus formation and halting infection.

### Identification of PaAvs5 Activator via TurboID Proximity Labelling

Based on previously known examples of the Avs family and the predicted protein-binding function of the TPR domain, we hypothesized that PaAvs5 is activated by a phage protein. To experimentally identify the phage protein responsible for activating PaAvs5-1’s enzymatic activity, we employed biotin proximity labeling^68^. To do this, we fused TurboID, a promiscuous biotin ligase, to the N-terminal of PaAvs5-1 and cloned this fusion construct into PAO1. This fusion retained the phage defense activity of PaAvs5-1 (Figure S1F), and we hypothesized that phage proteins interacting with PaAvs5 would show enriched biotinylation (Figure 5A).

**Figure 5.**
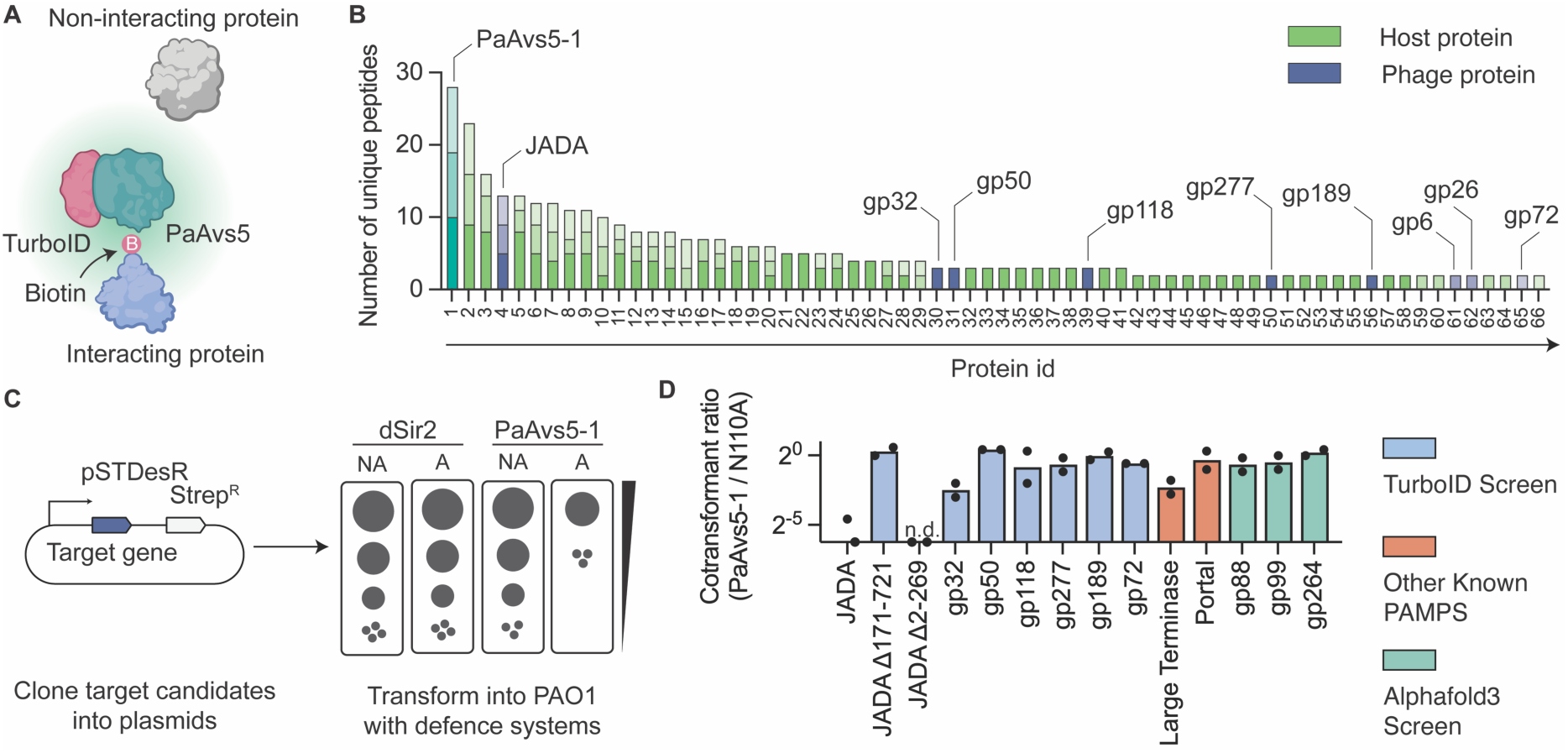
Identification and validation of phage proteins activating PaAvs5-1. (A) Schematic of the proximity biotinylation strategy using TurboID-tagged PaAvs5-1 to identify Pa36 phage proteins sensed by PaAvs5-1. Proteins in proximity to PaAvs5-1 are biotinylated by the TurboID tag; non-interacting proteins remain unmodified. (B) Mass spectrometry analysis of biotinylated proteins showing the number of unique peptides identified per protein in PAO1 expressing PaAvs5-1-TurboID. Host proteins are shown in green and Pa36 phage proteins in blue. PaAvs5-1 and detected phage proteins are annotated. Bars represent data from three biological replicates, with different transparency levels indicating individual replicates. Details of protein id are provided in Table S1. Raw shot gun proteomics data is available in the PRIDE database under project accession number PXD065050. (C) Schematic of the co-transformation assay. Candidate genes were cloned under a rhamnose-inducible promoter (pSTDesR plasmid; Streptomycin resistance) and transformed into PAO1 strains expressing either wild-type PaAvs5-1 or the catalytically inactive N110A mutant (pUCP20 plasmid (dSIR2); Carbenicillin resistance). (D) Transformant ratio (PaAvs5-1/N110A) shown in log scale for each candidate gene. Bars in purple represent candidates identified in the TurboID screen. Bars in orange indicate common phage-associated molecular patterns (PAMPs) known to be sensed by other defense systems. Activating proteins such as JADA and the large terminase show reduced transformation efficiency in cells expressing wild-type PaAvs5-1. Data represent ratios of mean values from three independent transformations. n.d. indicates “not detectable,” corresponding to zero colonies observed in transformation assays.

We found 66 unique biotinylated peptides from PAO1 cells carrying TurboID-PaAvs5-1 fusion when infected with Pa36 during our screening (Figure 5B). The most abundant unique peptide was derived from PaAvs5-1 which indicates self-biotinylation. Several abundant host proteins from PAO1 including phenylalanine-tRNA ligase alpha subunit, large ribosomal subunit proteins and DNA directed RNA polymerase subunits show enriched biotinylation. Nine Pa36 proteins were biotinylated of which only one protein, gp316, was enriched in all the three biological replicates of the experiment. This was a strong indication that PaAvs5-1 interacts with gp316, a protein of unknown function, and could potentially be the protein sensed by the system to activate its Sir2 catalytic activity.

To test whether these proteins were the activator of PaAvs5-1, we introduced a low copy pSTDesR plasmid encoding the phage proteins under a Rhamnose inducible promotor to PAO1 with and without the PaAvs5-1 system. Our rationale was that PaAvs5-1 activation by the phage target protein would diminish transformation efficiency since co-expression of both PaAvs5-1 and phage target protein would lead to the NAD^+^ hydrolysis and growth arrest (Figure 5C). PAO1 cells lacking PaAvs5-1 or carrying non-functional PaAvs5-1 mutants stably transformed with gp316. In contrast, transformants were obtained when PAO1 cells carried the dSir2 mutant of PaAvs5-1 and were transformed with gp316, but not when they carried the wild-type PaAvs5-1. This shows that gp316 activates the cell death function of PaAvs5-1. This verified that gp316 was one of the activators of PaAvs5-1 from our TurboID screen. We then successfully cloned six out of the eight remaining Pa36 proteins that showed enriched biotinylation. All six stably co-transformed with PaAvs5-1, and five of them showed similar transformation levels in both wild-type PaAvs5-1 and the dSir2 mutant (Figure 5D). gp32 also showed nearly a twofold reduction in transformation efficiency indicating that it could be a potential secondary activator.

We then tested if the identified activators can be sensed by PaAvs5 systems from the other clades. Surprisingly, we observed that gp316 was able to activate both PaAvs5-2 and PaAvs5-3 despite its broader anti-phage immunity profile (Figure S5A). Given its ability to activate all tested homologs of the PaAvs5 defense system, we renamed Pa36 gp316 as JADA (Jumbo phage Avs5 Defense Activator, Accession number: XCN26233.1).

We further attempted to identify the activators of PaAvs5-1 using AlphaFold2 and AlphaFold3-based protein–protein co-folding. Out of 356 jumbo phage proteins screened, three candidates (gp88, gp99, and gp264) showed co-folding with PaAvs5-1, with ipTM scores above 0.7 and low predicted alignment error, suggesting potential interaction (Figure S4). However, none of these proteins activated PaAvs5-1 in experimental assays (Figure 5D).

We also tested whether homologs of known AVAST system activators—specifically the large terminase subunit and portal proteins—could activate PaAvs5 systems. The large terminase subunit caused a twofold reduction in transformants in wild-type PaAvs5-1 and PaAvs5-3 compared to the PaAvs5-1 dSir2 mutant (Figure 5D, S5A). Similarly, the portal protein induced a modest reduction in transformants in the PaAvs5-2 system compared to the dSir2 control (Figure 5D, S5A). However, none of these candidates triggered an immune response as strong as that induced by JADA, which was the only activator to elicit a robust response across all three Avs5 clades. While we have not yet identified the activators from other phages, sequence homologs of gp316 were not found in the phages targeted by PaAvs5-2 and PaAvs5-3. This suggests that Avs5 systems can be triggered by multiple, distinct phage ligands. In summary, we identified an essential jumbo phage PAMP that can activate phylogenetically distant Avs5 systems.

### JADA is Potentially an Early Activator of PaAvs5

Since JADA activated all three PaAvs5 systems robustly, we next investigated its role in the jumbo phage lifecycle. Previous evidence suggests that JADA is essential for jumbo phage replication. Recent transcriptomics and proteomics data for phiKZ infection demonstrated that phiKZ_124, an ortholog of JADA is essential, highly expressed and is found among the most abundant phage proteins ten minutes post infection^69,70^. The abundance and early expression of JADA is consistent with the rapid initiation of NADase activity of PaAvs5-1 upon Pa36 infection.

To investigate the role of JADA, we first analyzed its distribution among all phage genomes. This revealed that this gene was primarily present in phage genomes already classified as *Chimalliviridae* (Figure S5B). We next checked the genomic context of JADA. This revealed that even in distant jumbo phages, JADA is invariantly located directly downstream of a non-virion DNA dependent RNA polymerase subunit (nvRNAP) and is transcribed in the same direction, suggesting coordinated expression (Figure 6A, S5C).

**Figure 6.**
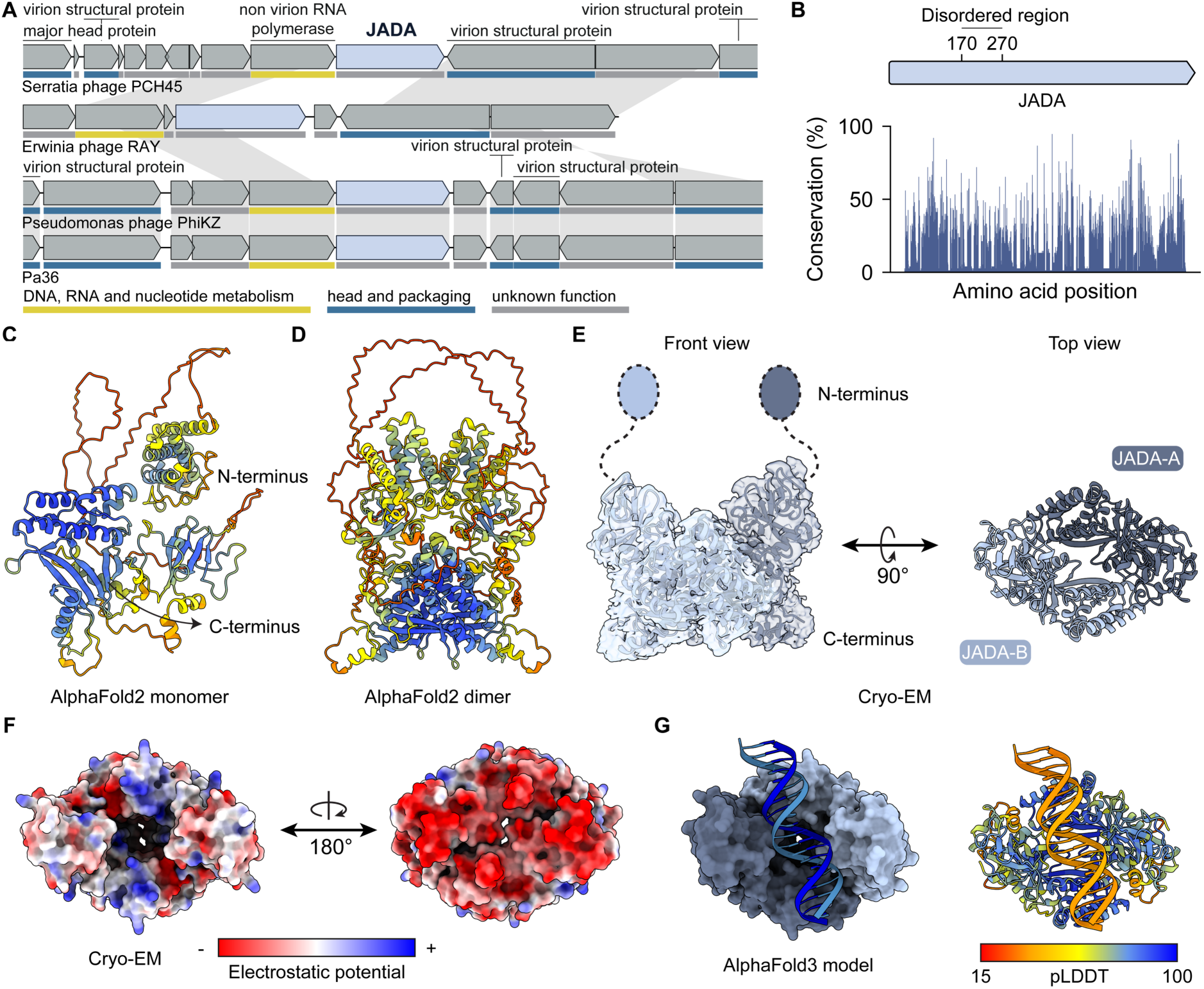
Structural characterization of JADA. (A) Genomic context of JADA showing its conserved position immediately downstream of a non-virion RNA polymerase subunit in the *Serratia* jumbo phage PCH45, the *Erwinia* jumbo phage RAY, *Pseudomonas* jumbo phages PhiKZ and Pa36. Hypothetical genes are not labelled. Only the first instance of each homolog is annotated, with homologous genes across genomes connected by grey lines. Putative gene functions are indicated by the color scheme shown below. (B) Amino acid conservation across JADA, based on an alignment of 75 homologs from diverse phages. Sequence conservation is shown as the percentage occurrence of the most frequent amino acid at each position. (C) AlphaFold2-predicted monomeric structure of JADA coloured by per-residue confidence (pLDDT; see G). (D) AlphaFold2-predicted dimeric structure of JADA, coloured by per-residue confidence (pLDDT, see G). (E) Cryo-EM reconstruction of the JADA complex at 2.1 Å resolution, shown in front and top views. The structure reveals a closed ring-like dimer, with disordered N-terminal extensions without detected electron density indicated as dotted lines. Individual monomer chains are labelled. Cryo-EM data is available in protein data bank under PDB id 9RP3. Cryo-EM refinement details are provided in Table S2 and Figure S6. (F) Electrostatic surface potential of the JADA from the cryo-EM structures. Two orientations (rotated 180°) reveal a strongly electronegative base (red).Z (G) AlphaFold3-predicted model of JADA bound to dsDNA. Left: surface representation of the JADA–DNA complex. Right: ribbon view of the complex coloured by pLDDT.

We attempted to characterize the interaction between PaAvs5-1 and JADA, but the PaAvs5-1 protein proved too unstable for *in vitro* biochemical assays or cryo-EM studies. Hence, we focused on understanding the structure and potential role of JADA. AlphaFold2 structural predictions (pTM = 0.53) show that JADA comprises an N-terminal domain (residues 1–170) connected to a C-terminal domain (residues 270– 721) by a long disordered or flexible region (Figure 6B, 6C, 6D). High-resolution cryo-EM (2.1 Å) of recombinantly purified JADA expressed in *E. coli* BL21-AI revealed a rigid C-terminal dimer (Figure 6E). The density map shows no evidence of bound ligands or ions within the central cavity. The base of the dimer is characterized by a highly negatively charged surface—features that may underlie function (Figure 6F). Additionally, no density was observed for the N-terminal region of the protein. Only the region corresponding to the C-terminal domain was solved by cryo-EM, consistent with the predicted disordered linker.

The N-terminal domain was well predicted in isolation (pLDDT > 90) and appears structurally stable but lacks significant Foldseek matches above a TM-score of 0.5, aside from hits to hypothetical proteins from related *Pseudomonas* phages in the BFVD database^52,71^. AlphaFold2 and AlphaFold3 models indicate that the N-terminal domain does not form homodimers (Figure 6D).

The C-terminal domain of JADA also lacks significant structural homologs, with Foldseek multimer failing to identify any confident hits in the PDB or BFMD databases (closest hit TM-score ∼0.3). The closest structural match for the full-length JADA monomer in AlphaFold DB has a TM-score of 0.57, again corresponding to a related *Pseudomonas* hypothetical phage protein. Functional annotation of JADA via ProteInfer suggests a potential DNA-binding and regulatory role^72^. Consistent with this function, AlphaFold3 modelling of the JADA C-terminal allows plausible DNA docking within a positively charged surface cavity of the homodimer (Figure 6G).

To determine which region of JADA activates the PaAvs5 system, we generated domain deletion constructs within the same pSTDesR plasmid: one encoding only the N-terminal domain (JADA Δ270–721) and another encoding only the C-terminal domain (JADA Δ2–269). When expressed alone, the N-terminal domain failed to activate PaAvs5 (Figure 5D). In contrast, co-expression of the C-terminal domain with inactive PaAvs5 yielded very few transformants under standard induction conditions, suggesting that the C-terminal domain may be toxic even in the absence of PaAvs5 activity. We then repeated the transformation assay under non-induced conditions, relying on leaky plasmid expression. Under these conditions, we recovered transformants with the inactive PaAvs5 system but not with the active PaAvs5 from all three clades (Figure 5D). These results indicate that the C-terminal domain of JADA is sufficient to activate Avs5 systems from all three clades.

## Discussion

Among the various strategies in the ongoing bacteria–phage arms race, the sequential genome compartmentalization employed by *Chimalliviridae* phages (nucleus forming jumbo phages) is particularly unique. This highly regulated lifecycle through sequential compartmentalization of phage genome allows them to evade all known DNA-targeting defense systems^1,2,4,6^. In response, bacteria have evolved alternative, specialized strategies that bypass DNA targeting altogether, instead recognizing phage-derived transcripts or proteins to trigger defense responses^25–27^. Here, we demonstrate that AVAST Type V systems from *P. aeruginosa* (PaAvs5), spanning three distinct clades, display a conserved robust defense against nucleus-forming jumbo phage (Figure 7). PaAvs5 is a member of the STAND superfamily, an ancient and widely distributed class of signal transduction proteins found across all domains of life.

**Figure 7.**
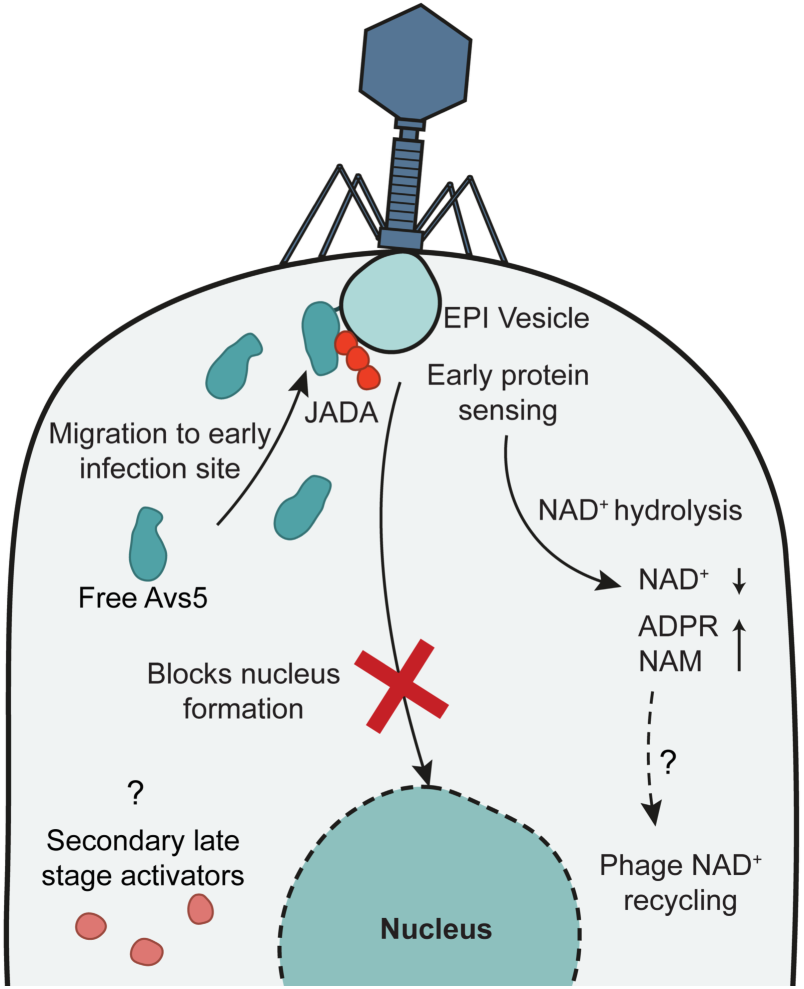
Model for Avs5-mediated defense against jumbo phages. Following infection, Avs5 migrates to the early infection site where it senses the early-expressed phage protein JADA. This sensing activates NAD^+^ hydrolysis by Avs5, leading to a transient depletion of cellular NAD^+^ levels. Although phage-encoded NAD^+^ recycling pathways may restore balance, the initial disruption is sufficient to block phage nucleus formation and thereby impair phage propagation. Dashed arrows indicate hypothetical mechanisms such as evasion strategies and the involvement of potential secondary late-stage activators.

The nucleus forming ability of jumbo phages is severely impacted by PaAvs5. In most cells containing PaAvs5, jumbo phage infection is halted at the EPI vesicle stage. A smaller group, about 10%, goes on to develop a phage nucleus after 90 minutes, doubling the time needed to reach this step. Such strong response might be mediated by rapid sensing and migration of PaAvs5 to infection sites. In nearly 75% of infected cells, PaAvs5 forms multiple distinct foci that colocalize with the EPI vesicle. This suggests PaAvs5 is recruited to this early transcription site where polysomes emerge from the vesicle surface. What makes PaAvs5 unique is that foci are not coupled to sensing alone, as inactive sensor mutant, Sir2 mutant and ATPase mutants were able to form such foci.

Upon sensing, PaAvs5-1 triggers a rapid drop in NAD^+^ levels due to strong hydrolysis within ten minutes of infection, consistent with activation by an early-stage phage factor. NAD⁺ and its hydrolysis product ADPR return to near-basal levels within the next ten minutes. While the mechanism remains unclear, the phage may encode NAD^+^ recycling enzymes that convert ADPR back to NAD^+73,74^. Despite the transient nature of this response, it is sufficient to protect the host, as the bacterial genome remains intact.

PaAvs5 systems, though exhibiting diverse anti-phage activity profiles and low sequence similarity, converge on sensing the same early-expressed phage protein to initiate an immune response. We identified and validated this shared trigger—an essential protein we named JADA (Jumbo phage Avs5 Defense Activator). Remarkably, JADA activates PaAvs5 systems across all three phylogenetic clades, enabling a robust defense that suppresses phage plaques by nearly seven orders of magnitude. High-resolution cryo-EM of purified JADA homodimer revealed a novel protein fold with no identifiable sequence or structural homologs outside of related jumbo phages. JADA is composed of two structural domains, with the larger C-terminal domain responsible for PaAvs5 activation. Furthermore, its conserved genomic positioning next to the non-virion RNA polymerase and gene ontology predictions point to a possible role in transcription during early phage infection. Together with the observed colocalization of PaAvs5 with the EPI vesicle, these findings suggest that JADA may function in early phage protein expression.

In addition to JADA, we identified two additional phage-encoded proteins that may activate PaAvs5 systems, albeit with much lower potency. In the case of PaAvs5-1, both the large terminase subunit and a hypothetical protein (gp32) from Pa36 induced moderate toxicity, indicative of system activation. The large terminase subunit also triggered PaAvs5-3 to a similar extent but failed to activate PaAvs5-2. Similarly, the portal protein activated the PaAvs5-2 system to similar levels. These findings support a model in which Avs5 systems incorporate layered sensing mechanisms, with highly conserved late-stage triggers potentially serving as fail-safes if the primary and conserved early-stage trigger is absent or evaded. Notably, the broad-spectrum activity of PaAvs5-3 against three unrelated phages—a jumbo phage, an RNA phage, and a siphophage—further suggests the existence of additional, unidentified triggers. These findings, together with recent results from CapRel^75^ and Avs2^35^, suggest that protein sensing defense systems may respond to more than one phage protein trigger, rather than a single specific target as previously thought.

It is becoming increasingly clear that bacteria have evolved dedicated strategies to counter jumbo phages, despite the sophisticated mechanisms these phages use to evade host defenses. The recently characterized Juk system directly target the EPI vesicle, an early-stage evasion structure employed by jumbo phages^27^. Our findings now show that Avs5 systems respond to components emerging from the EPI vesicle. Together, these systems illustrate a broader strategy in which conserved phage countermeasures are used as vulnerabilities by bacterial immune defenses. It is likely that future studies will uncover additional defense systems specialized to target distinct stages of the jumbo phage lifecycle.

## Supporting information

Supplementary document containing Figure S1-S6 and Table S2-S4

## Acknowledgments

This work was supported by grant from the European Research Council (ERC) CoG under the European Union’s Horizon 2020 research and innovation program (grant agreement no. 101003229) awarded to S.J.J.B. A.M. is supported by grants from Koningin Wilhelmina Fonds (KWF, grant agreement no. 15602) and Nederlandse Organisatie voor Wetenschappelijk Onderzoek (NWO, grant agreement no. OCENW.XS23.1.006). We acknowledge the infrastructure provided by Kavli Nanolab Imaging Centre for optical microscopy. M.P. was supported by the Peter und Traudl Engelhorn Stiftung. B.E.C. was supported by the Swiss National Science Foundation. We thank Alexander Myasnikov, Bertrand Beckert, and Sergey Nazarov (Dubochet Center for Imaging, EPFL-UNIL-UNIGE, Switzerland) for assistance with cryo-EM data collection. We thank Prof. Rob Lavigne (KU Leuven) for kindly providing plasmid pSTDesR.

## Author contributions

Conceptualization, A.M., S.J.J.B.; Methodology, A.M., S.J.J.B.; Formal Analysis, A.M., A.R.C., D.F.v.d.B., M.P. (Marting Pacesa); Investigation, A.M., A.R.C., D.F., D.F.v.d.B., H.v.d.B., A. Z-D., M.Pt. (Martin Pabst), M.P.; Visualization, A.M., D.F.v.d.B., H.v.d.B., M.P.; Writing – Original Draft, A.M.; Writing – Review & Editing, A.M., A.R.C., M.P., S.J.J.B.; Resources, A.M., M.P., B.E.C, S.J.J.B.; Supervision, A.M., S.J.J.B; Funding Acquisition, A.M., S.J.J.B.

## Materials and Methods

### Bacteria and Phages

Three clinical isolates of *Pseudomonas aeruginosa* (accession number: GCA_030069065.1, MCO2951615.1, MCO2344054.1) obtained from the University Medical Center Utrecht was utilized to amplify the PaAvs5 defense systems using PCR with suitable primers. For cloning experiments involving the PaAvs5 systems into plasmid pUCP20, *Escherichia coli* strain DH5α and *Pseudomonas aeruginosa* strain PAO1 were employed. All bacterial cultures were maintained in Lysogeny Broth (LB) at 37°C with shaking at 180 RPM or on LB agar (LBA, 1.5% agar w/v) plates at 37°C, unless specified otherwise. Strains containing plasmid pUCP20 were grown in LB media supplemented with either 100 µg/mL ampicillin (for *E. coli*) or 200 µg/mL carbenicillin (for *P. aeruginosa*).

The phages used in this study, except for *Pseudomonas* phage PP7 (obtained from LGC Standards), were obtained from the Fagenbank. Phages were propagated in liquid cultures with PAO1, followed by centrifugation at 3,000 × g for 15 minutes, filtration through 0.2 µm PES filters, and storage as lysates at 4°C until needed.

### Phylogenetic Tree of Avs5

The phylogenetic tree of Avs5 was built with the Avs5 protein sequences provided by Gao et al. (2020)^30^. These sequences were aligned using Muscle v5.1.0^76^ (default settings) and trimmed using trimAl v1.5.0 (default settings). The resulting trimmed alignment was used to build and bootstrap a phylogenetic tree using IQ-Tree2 v2.3.6^77^ (-B 1000, --alrt 1000, -m TEST). This phylogenetic tree was visualized using iTol v5.0^78^.

### Annotation of Functional Domains

CDD v3.21^79,80^ and InterProScan v5^81^ were used to annotate the functional domains. Additionally, protein tandem repeat domains were searched for using TPRpred v1.0^82^ and HHrepID v1.0^83^ to detect TPR and de-novo tandem repeats, respectively. As well as a protein sequence and structural comparison using blastp v2.16.0^84^ all-versus-all and AlphaFold3^85,86^.

### Cloning of PaAvs5 Systems into PAO1

The PaAvs5 systems and their promoter regions were amplified from *Pseudomonas aeruginosa* strains using the primers (Integrated DNA Technologies) listed in the Table S3 and Q5 High-Fidelity DNA Polymerase (New England BioLabs) supplemented with Q5 High GC Enhancer (New England BioLabs). The resulting PCR products mixed with loading dye were visualized on 1% agarose gels supplemented with SYBR Safe DNA Gel Stain (Invitrogen), and the target bands were excised and purified with the Zymoclean Gel DNA Recovery Kit (Zymo Research). Plasmid pUCP20 was digested with BamHI and EcoRI (New England BioLabs), treated with FastAP (Thermo Scientific) to remove any overhangs, and then purified using the Zymo DNA Clean & Concentrator Kit (Zymo Research). The amplified PaAvs5 systems were then cloned into a digested pUCP20 plasmid using the NEBuilder HiFi DNA Assembly Master Mix (New England BioLabs) and subsequently transformed into high-efficiency NEB 5-alpha competent *E. coli* (New England BioLabs), following the manufacturer’s protocol. After growing the transformed *E. coli* in S.O.C. (Super Optimal broth with Catabolite repression) for 1 hour at 37°C, the culture is plated on LB agar supplemented with 100 µg/mL ampicillin and incubated overnight at 37°C.

Plasmids were extracted using the GeneJet Plasmid Miniprep Kit from overnight cultures from individual colonies from the transformed plate and verified through nanopore sequencing (Macrogen). Confirmed plasmids were introduced into *P. aeruginosa* strain PAO1 via electroporation. To prepare electrocompetent PAO1 cells, an overnight culture of PAO1 was centrifuged at 16,000 × g for two minutes at room temperature^87^. The pellet was washed twice with 300 mM sucrose and resuspended in the same solution. For electroporation, 200–500 ng of plasmid DNA was added to 20 µl of the electrocompetent PAO1 suspension. The mixture was transferred to a 2.5 mm electroporation cuvette and electroporated at 2.5 kV. The transformants were then plated on LB agar supplemented with 200 µg/mL carbenicillin.

Point mutations in the Sir2 or ATPase domains, as well as deletions within the sensor or putative signal peptide domains of Avs5 in pUCP20, were introduced using round-the-horn site-directed mutagenesis with 5’ phosphorylated primers (Integrated DNA Technologies) and Q5 DNA Polymerase supplemented with Q5 High GC Enhancer (New England BioLabs). The resulting PCR products were digested with DpnI (New England BioLabs), separated on 1% agarose gels, and the bands were excised and purified using the Zymo Gel DNA Recovery Kit. The amplified plasmids were then ligated using T4 DNA ligase (New England BioLabs) at room temperature for two hours and transformed into chemically competent NEB 5-alpha Competent *E. coli* according to the manufacturer’s instructions.

Plasmids were extracted using the GeneJET Plasmid Miniprep Kit and verified by nanopore or Sanger sequencing (Macrogen). The confirmed plasmids were subsequently introduced into *P. aeruginosa* strain PAO1 via electroporation as described above. The transformants were then plated on LB agar supplemented with 200 µg/mL carbenicillin.

### Cloning of PaAvs5-1 Fusion Proteins into PAO1

The gBlocks encoding mNeonGreen and TurboID (Integrated DNA Technologies) were amplified using primers with overhangs homologous to the PaAvs5-1 gene and the pUCP20 plasmid. The primer sequences are listed in Table S3. The amplified PaAvs5-1 constructs were cloned into the digested pUCP20 plasmid using the NEBuilder HiFi DNA Assembly Master Mix (New England BioLabs) and subsequently transformed into high-efficiency NEB 5-alpha competent *E. coli* (New England BioLabs). The transformed *E. coli* is plated on LB agar supplemented with 100 µg/mL ampicillin.

Plasmids were extracted using the GeneJET Plasmid Miniprep Kit and verified by sequencing (Macrogen). The confirmed plasmids were then introduced into PAO1 by electroporation as described above. The transformants were then plated on LB agar supplemented with 200 µg/mL carbenicillin.

For both mNeonGreen and TurboID, plasmids were constructed with the fusion at either the amino or carboxy terminus of PaAvs5-1. The effect of the fusion on PaAvs5 activity was evaluated using an efficiency-of-plating assay with the phage Pa36. Experiments were subsequently conducted with the construct that had minimal impact on PaAvs5 system activity.

### Cloning of Phage Genes

Phage genes were PCR-amplified using Q5 DNA polymerase (New England BioLabs) supplemented with Q5 High GC Enhancer (New England BioLabs) with primers (Integrated DNA Technologies) containing overhangs homologous to the pSTDesR plasmid^88^. Primer sequences are listed in Table S3. The amplified phage gene constructs were cloned into the PCR-amplified pSTDesR plasmid using NEBuilder HiFi DNA Assembly Master Mix (New England BioLabs) and subsequently transformed into high-efficiency NEB 5-alpha competent *E. coli* (New England BioLabs). The phage gene inserts are under the control of a rhamnose-inducible promoter, pRhaBAD. Transformed *E. coli* cells were plated on LB agar supplemented with 12.5 µg/mL streptomycin.

Plasmids were extracted using the GeneJET Plasmid Miniprep Kit and verified by sequencing (Macrogen). The confirmed plasmids were then introduced into PAO1 by electroporation as described above. The transformants were then plated on LB agar supplemented with 25 µg/mL streptomycin.

### Efficiency of Plating

Phage stocks were diluted 10-fold serially in LB in a 96-well plate, and the resulting dilutions were spotted onto double-layer agar plates supplemented with 200 µg/mL carbenicillin and inoculated with *P. aeruginosa* PAO1. These plates contained either the empty pUCP20 plasmid, pUCP20 with PaAvs5-1, mutant PaAvs5-1, or PaAvs5-1 with a fusion protein, following the small plaque drop assay method. To assess the anti-phage activity of the defense systems, the fold reduction in phage infectivity was calculated by comparing the phage infectivity in the PAO1 strain carrying the defense systems encoded by pUCP20 to that in the strain carrying the empty plasmid.

### Liquid Culture Collapse Assays

Overnight bacterial cultures were diluted to an OD_600_ of approximately 0.1 in LB and transferred into 96-well plates. The bacterial strains containing pUCP20 plasmids were supplemented with 200 µg/mL carbenicillin. Phages were added at multiplicities of infection (MOIs) of 0.01 and 10, and the plates were incubated at 37°C in an Epoch2 microplate spectrophotometer (Biotek) with double orbital shaking. OD_600_ measurements were taken every 10 minutes for 24 hours to monitor bacterial growth.

### Cell Lysate Preparation for NAD^+^ and ADPR LC-MS

Overnight cultures of *P. aeruginosa* PAO1 strains containing the pUCP20 plasmid with PaAvs5-1, the PaAvs5-1 N110A point mutant, or no defense systems were diluted 1:100 in 250 mL LB and incubated at 37°C with shaking at 180 RPM. When the OD_600_ reached 0.3, 50 mL of uninfected culture (corresponding to time = 0 minutes) was removed. Phage stock was then added to the remaining culture to achieve a MOI of 3. The flasks were incubated at 37°C with shaking at 180 RPM for the duration of the experiment.

At each time point, 50 mL of the culture was removed. Immediately after sampling, the tubes were centrifuged at 4 °C for five minutes at 3900 RPM to stop the phage infection process. The supernatant was discarded, and the pellet was frozen at -20 °C. To extract metabolites and lysing the cells, 1 mL of 100 mM phosphate buffer (pH 8) consisting of Potassium Phosphate Dibasic (∼ 93.5 mM) and Potassium Phosphate Monobasic (∼ 6.5 mM) and supplemented with 4 mg/mL lysozyme was added to each pellet. The tubes were thawed on ice for ten minutes and then transferred to 2 mL FastPrep tubes (MP biomedicals) and 0.1 g of cell disruption media composed of 0.1 mm glass beads (Scientific Industries Inc) were added. To each tube, 0.5 µmol of β-Nicotinamide Adenine Dinucleotide Sodium Salt (Sigma) and Adenosine 5’- Diphosphoribose Sodium (Sigma) were added as internal standards. The cells were then lysed using the FastPrep-25 5G bead beater (MP biomedicals) for 40 seconds at the manufacturer’s recommended settings for *E. coli*. The tubes were centrifuged at 4 °C for ten minutes at 15,000 × g to separate the beads from the lysate. The supernatant was then transferred to an Amicon Ultra-0.5 centrifugal filter unit (3 kDa) and centrifuged for one hour at 4°C at 6000 × g. The filtrate was collected, frozen at - 20°C and subsequently used for LC-MS analysis.

### Quantification of NAD± and ADPR by LC-MS

The LC-MS analysis was performed using an Agilent LC/MS system consisting of a high-pressure liquid chromatography set-up coupled to a triple-quadrupole (QQQ) mass spectrometer (G6460C) equipped with a standard electrospray ionization (ESI) source. Both systems were operated through MassHunter data acquisition software (version 10.1). 1 µL of each sample was injected into the column of the HPLC. NAD^+^ and ADPR were delivered to a CSH C18 guard column and a CSH C18 column (Waters) (2.1 mm by 50 mm, 1.7-µm pore size) at 30°C with a flow rate of 0.3 mL/min using the following binary gradient: 0% B (ACN, 25 mM FA), ramp to 85% B in 9 min, followed by a 30 sec hold at 85% B, a 2 min ramp back to 0% B, and a 3 min re-equilibration (A, 20 mM ammonium formate). Next, the metabolites were eluted from the column, and the eluent was sprayed into the mass spectrometer, which was operated in data-dependent mode, specifically in dynamic multiple-reaction monitoring (dMRM) mode using transitions. Each MRM transition was generated by optimizing the fragmentor voltage and the collision energy. The dMRM was acquired in positive mode with a cycle time of 500 ms. Data processing was done using Skyline^89^.

For the quantification of NAD^+^ and ADPR, a calibration curve (0–500 μM) using the standard addition method was employed. The batch design for running the samples was as follows: first, the calibration curve was analysed, followed by a blank sample. Then, the randomized samples were run, with a blank injected after every five samples to monitor carry-over. Subsequently, the calibration curve was injected again.

In Skyline, the peaks corresponding to NAD^+^ and ADPR were integrated for quantification, and the areas under the curves were exported for further analysis. A linear calibration curve (R^2^ > 0.9) was obtained for concentration calculations of each compound in all the samples. The measured concentrations are multiplied by 50 to account for the lysate preparation step, where 50 mL was concentrated to 1 mL.

### Confocal Fluorescence Microscopy of PaAvs5-1

Exponentially growing PAO1 cultures (OD_600_ ≈ 0.3) containing either the fluorescently tagged PaAvs5-1 or negative were infected with phage at an MOI ≥3 so that all bacteria are infected. The phage was allowed to adsorb for 10 minutes at 37°C. The cells were centrifuged at 9,000g for 1 minute. The cell pellet was resuspended in 100 μL of LB. 2 μL each of 4’,6-Diamidino-2-Phenylindole (DAPI, 5 mg/mL) and MitoTracker Deep Red FM (5 mg/mL) were added to the cell suspension. The resuspended cells (2 μL) were spotted onto 1% agarose pads.

Visualization was performed using a Nikon A1R/SIM laser scanning confocal microscope (inverted Nikon Ti Eclipse body) equipped with a 100× oil immersion objective (SR Apo TIRF; numerical aperture 1.49). Lasers were used sequentially to excite the different channels, from longest to shortest wavelength: MitoTracker Deep Red FM with 640 nm (emission filter: 700/75 nm), mNeonGreen with 488 nm (emission filter: 525/50 nm), and DAPI with 405 nm (emission filter: 450/50 nm), all passed through a 405/488/543/640 excitation dichroic. Z-stacks were acquired using a Nikon A1 Piezo Z Drive at intervals of either 0.2 or 0.1 µm (10–20 slices), capturing different planes of all bacteria within the field of view (512 × 512 pixels, corresponding to 36.79 × 36.79 µm, satisfying Nyquist criteria). A pinhole size corresponding to 1.2 AU referenced to the shortest wavelength was used. Images were acquired at 12-bit depth using a Galvano scanner with Nikon NIS-Elements software. Image analysis was performed using Fiji, and Napari. To perform image analysis, bacteria were segmented using Cellpose^90^ using Deepbacs^91^ model in Napari.

### Proximity Labelling to Identify PaAvs5 Activator

The TurboID-PaAvs5 construct was cloned into the pUCP20 plasmid and introduced into PAO1 cells. The PaAvs5 system was N-terminally tagged with TurboID, as this configuration retained its defense phenotype against phage Pa36. In contrast, C-terminal tagging abolished the defense activity. Cultures were grown overnight and subsequently diluted to an OD_600_ of 0.1 in fresh medium supplemented with 500 µM biotin. Upon reaching an OD_600_ of 0.3, cells were infected with phage Pa36 at a MOI of 3.

### Cell Harvesting, Protein Extraction, On Beads Enrichment and Proteolytic Digestion

At 20 minutes post-infection, cultures were harvested by centrifugation at 9000 g for 10 minutes at 4 °C. Cell pellets were stored at −80 °C until further processing.

For protein extraction, pellets were resuspended in lysis buffer (100 mM Tris-HCl pH 7.5, 150 mM NaCl, 5% glycerol, 1 mM DTT) and lysed using a sonication. Cell debris and beads were removed by centrifugation, and only the supernatant was retained. Biotinylated proteins were enriched via affinity purification using Strep-Tactin XT beads. Beads were washed three times with lysis buffer and retained for on-bead processing.

Enriched proteins were reduced with 10 mM dithiothreitol (DTT) in 100 mM ammonium bicarbonate (ABC) for 60 minutes at 37 °C with shaking at 300 rpm, followed by alkylation with 20 mM iodoacetamide (IAA) in 100 mM ABC for 30 minutes at room temperature in the dark. Proteins were then digested on-bead overnight (∼18 hours) at 37 °C with Trypsin Sequencing Grade, also at 300 rpm.

Following digestion, beads were removed by centrifugation, and the peptide-containing supernatant was subjected to solid-phase extraction using an Oasis HLB 96-well µElution Plate. The plate was conditioned with methanol, equilibrated with LC-MS grade water, and samples were loaded and washed twice with 5% methanol. Peptides were eluted sequentially with 200 µL of 2% formic acid in 80% methanol followed by 200 µL 1 mM ABC in 80% methanol. The combined eluates were dried using a SpeedVac evaporator at 45 °C for 3–4 hours.

Prior to LC-MS analysis, samples were resuspended in 15 µL of 3% acetonitrile + 0.01% trifluoroacetic acid (TFA) in LC-MS grade water and analyzed using a nanoLC-Q Exactive Plus Orbitrap system.

### Shotgun Proteomic Analysis

An aliquot of each sample was analyzed using a nano-liquid-chromatography system consisting of an EASY nano LC 1200 equipped with an Acclaim PepMap RSLC RP C18 reverse phase column (75 mm x 150 mm, 2 mm) coupled to a QE plus Orbitrap mass spectrometer (Thermo Scientific, Germany). Solvent A was H_2_O containing 0.1% formic acid, and solvent B consisted of 80% acetonitrile in H_2_O, containing 0.1% formic acid. The flow rate was maintained at 350 nL/min. The Orbitrap was operated in top 10 data dependent acquisition (DDA) mode, acquiring peptide signals form 350–1250 m/z, at 70K resolution in MS1 with an AGC target of 3e6 and max IT of 100 ms. An aliquot of approx. 100 ng protein digest was analyzed using a linear gradient from 2 to 40% B over 60 minutes. MS2 acquisition was performed at 17.5K resolution, with an AGC target of 5e5, and a max IT of 100 ms, using a NCE of 28. Unassigned, singly charged as well as >4 charged mass peaks were excluded.

### Database Searching of Mass Spectrometric Raw Data

Mass spectrometric raw data were processed using PEAKS Studio X (Bioinformatics Solutions Inc., Canada) for database searching and de novo sequencing. Database searching was performed using the reference proteome of Pseudomonas aeruginosa (UP000002438, strain ATCC 15692) including phage sequences, and the cRAP proteome (https://www.thegpm.org/crap/), allowing 20 ppm parent ion and 0.02 Da fragment mass error and up to 3 missed cleavages. Carbamidomethylation was set as fixed and Biotinylation (+ 226.08 Da) as variable modification. Database search further used decoy fusion for estimation of false discovery rates (FDR) and subsequent filtering of peptide spectrum matches for 1% FDR, and 2 unique peptides per protein.

### Validation of Activators of PaAvs5

Overnight cultures of P. aeruginosa PAO1 strains harboring either wild-type PaAvs5-1 or the catalytically inactive dSir2 mutant (N110A), both expressed from the pUCP20 plasmid, were grown and made electrocompetent as described above. Candidate phage genes, cloned into pSTDesR plasmids under the control of a rhamnose-inducible promoter (100 ng/μL), were electroporated into these electrocompetent cells using the same electroporation protocol.

Following transformation, 1 mL of LB medium was added, and cells were incubated at 37 °C for 1 hour. Cultures were then centrifuged, and the pellets were resuspended in 200 μL of LB. Each transformation was tested on both induced and uninduced conditions by spotting twofold serial dilutions (5 μL) onto LB agar plates containing carbenicillin and streptomycin, either without inducer or supplemented with 2.5 mM rhamnose. Each transformation was performed in duplicate. Plates were allowed to dry, then incubated overnight at 37 °C. Colony-forming units were counted the next day, and activation was assessed by comparing colony numbers in the wild-type versus dSir2 mutant.

### JADA Locus Visualization

We downloaded the genomes of bacteriophages classified under the family Chimalliviridae as described in Prichard et al. (2023)^92^. Genome annotation was performed using Phannotate^93^, Pharroka^94^, and Phold. To identify homologs of the JADA protein, we constructed a hidden Markov model (HMM) profile based on PSI-BLAST^95^ hits with >60% query coverage. Sequences were aligned using Muscle v5.1.0^76^, and the HMM profile was built using hmmbuild^96^. The genomic locus of JADA homologs was visualized using LoVis4u^97^.

### JADA Protein Purification

The JADA gene (Pa36 gp316) was amplified from *Pseudomonas aeruginosa* phage Pa36 genomic DNA using Q5 polymerase (NEB) with primers designed to incorporate homology arms matching the pACYCDuet-1 vector. The PCR product was cloned into the vector via Gibson assembly, placing a N-terminal 6×His tag for affinity purification. The resulting plasmid was confirmed by nanopore sequencing and transformed into E. coli BL21-AI cells. They were then plated on chloramphenicol-containing LB agar plates.

For expression, these cells were inoculated in LB medium supplemented with chloramphenicol and grown at 37 °C until the culture reached an OD_600_ of 0.6. Cultures were then cold-shocked on ice for 1 hour and induced with 1 mM IPTG and 0.2% arabinose. Following induction, cells were transferred to 20 °C and incubated overnight with shaking.

Cells were harvested by centrifugation using an Avanti J-26 XP centrifuge at 3900 × g for 30 minutes at 4 °C. The supernatant was discarded, and the cell pellet was resuspended in ice-cold PBS. This suspension was centrifuged again under the same conditions, and the supernatant was removed. The resulting washed pellets were stored at –80 °C until further processing.

Pellets were thawed on ice and resuspended in lysis buffer (100 mM Tris-HCl pH 7.5, 300 mM NaCl, 1 mM DTT, 5% glycerol) supplemented with cOmplete protease inhibitor cocktail (Roche). Cell lysis was performed at 1 bar using a continuous flow cell disruptor (Constant Systems), and lysates were clarified by centrifugation in an Avanti centrifuge at 16000 × g for 30 minutes at 4 °C. The resulting supernatant was filtered through a 0.45 μm syringe filter prior to affinity purification.

For affinity purification, the lysate was loaded onto a Ni-NTA His-Select column (pre-equilibrated with ice cold lysis buffer supplemented with 25 mM imidazole). The column was washed with 10 column volumes of the same buffer, and the bound protein was eluted with lysis buffer containing 250 mM imidazole. Eluted fractions were concentrated and further purified by size exclusion chromatography using a Superdex 200 Increase 10/300 GL column on an ÄKTA system, with lysis buffer as the running buffer. The purified JADA protein was snap-frozen in liquid nitrogen and sent for cryo-EM analysis.

Prior to cryo-EM, thawed proteins (JADA alone, PaAvs5-1 alone, or mixture) were purified once more by size exclusion chromatography using a Superose 6 Increase column.

### JADA Cryo-EM Structure Determination

Purified JADA protein was applied to a glow-discharged 300-mesh holey carbon grid (Au 1.2/1.3, Quantifoil Micro Tools), blotted for 4 s at 95% humidity, 10 °C, plunge-frozen in liquid ethane (Vitrobot, Thermo Fisher Scientific) and stored in liquid nitrogen. Data collection was performed with automation program EPU (Thermo Fisher Scientific, v.2.12.1) on a 300 kV FEI Titan Krios G4 microscope equipped with a FEI Falcon IV detector. A total of 15624 micrographs were recorded with a pixel size of 0.726 Å. Acquired cryo-EM data was processed using cryoSPARC (v.4.6.2)^98^. Gain-corrected micrographs were imported, and micrographs with a resolution estimation worse than 6 Å were discarded after patch contrast transfer function estimation. Initial particles were picked using a blob picker with 70-105 Å particle size. Particles were extracted with a box size of 280 × 280 pixels, down sampled to 100 × 100. After two-dimensional classification, clean particles were used for ab initio three-dimensional reconstruction. After several rounds of three-dimensional classification, the class with most detailed features was reextracted using full box size and subjected to non-uniform and local refinement to generate high-resolution reconstructions. The local resolution was calculated and visualized using ChimeraX (v.1.9, UCSF).

For structure building, we used an initial model generated by ModelAngelo^99^, which was then used as template input for ColabFold^100^ reprediction. Full dimer model was then fitted into density and manually refined using Coot (v0.9.5)^101^ Atomic model refinement was performed using Phenix.real_space_refine (v.1.20.1-4487)^102^. The quality of the refined model was assessed using MolProbity (v.4.5.1)^103^. The refined atomic models and corresponding cryo-EM maps were deposited under PDB accession code 9RP3 and EMDB accession code EMD-54139. Details of data collection and refinement statistics are shown in supplementary Table S2.

### PaAvs5-1 Protein Purification

Expression and purification of PaAvs5-1 were performed using procedures similar to those described above for JADA. The gene was amplified with homology to the p13SS expression vector and cloned via Gibson assembly. The construct included a N terminal Twin-Strep-SUMO tag for affinity purification and was expressed in *E. coli* BL21-AI using streptomycin selection. Additional variants were also tested: (1) a codon-optimized version of PaAvs5-1 for *E. coli* expression; (2) a version cloned with a N-terminal 6×His tag in the pACYCDuet-1 backbone (as used for JADA); and (3) an N110A point mutant of PaAvs5-1.

For Twin-Strep–SUMO-tagged constructs, purification was carried out using StrepTactin resin. After cell lysis (see above for buffer and disruption method), the cleared lysate was passed over StrepTactin beads, washed with lysis buffer (100 mM Tris-HCl pH 7.5, 300 mM NaCl, 1 mM DTT, 5% glycerol), and eluted in the presence of 50 mM biotin. Expression was verified using SDS-PAGE and then purified further using size exclusion chromatography as stated above.

Despite testing multiple expression constructs and purification strategies—including the addition of ATP or the non-hydrolysable analog AMP-PNP—PaAvs5 consistently yielded very low amounts of protein. Attempts to visualize particles by cryo-EM, either for PaAvs5 alone or in combination with JADA, failed to produce any visible particles.

## Supplemental information

**Document S1.** Figures S1-S6, Table S2-S4

**Table S1** Excel Sheet of Protein IDs and Annotations for TurboID Interactome Screen, Related to Figure 5B.

