## Supplementary document containing Figure S1-S6 and Table S2-S4 for "Molecular basis for anti-jumbo phage immunity by AVAST Type 5"

### List of contents

**Figure S1** Sequence Conservation and Functional Analysis of Avs5 Domains and Motifs Related to Figure 1.

**Figure S2** Fluorescence Microscopy of PaAvs5-1 and Domain Mutants, Related to Figure 4.

**Figure S3:** Localization of PaAvs5-1 During Pa34 Infection, Related to Figure 4.

**Figure S4** AlphaFold3 Cofolding Predictions to Identify Potential Interactors of PaAvs5-1, Related to Figure 5

**Figure S5** Conservation and Genomic Context of JADA Homologs Across Jumbo Phages, Related to Figure 6.

**Figure S6** Cryo-EM Map Quality and Resolution Estimation of JADA Homodimer, Related to Figure 6.

**Table S2** Cryo-EM Data Collection, Processing, and Model Validation Statistics for JADA/Gp316 Homodimer (PDB: 9RP3, EMDB: 54139), Related to Figure 6.

**Table S3:** List of Primers Used in This Study.

**Table S4:** List of Plasmids Used in This Study

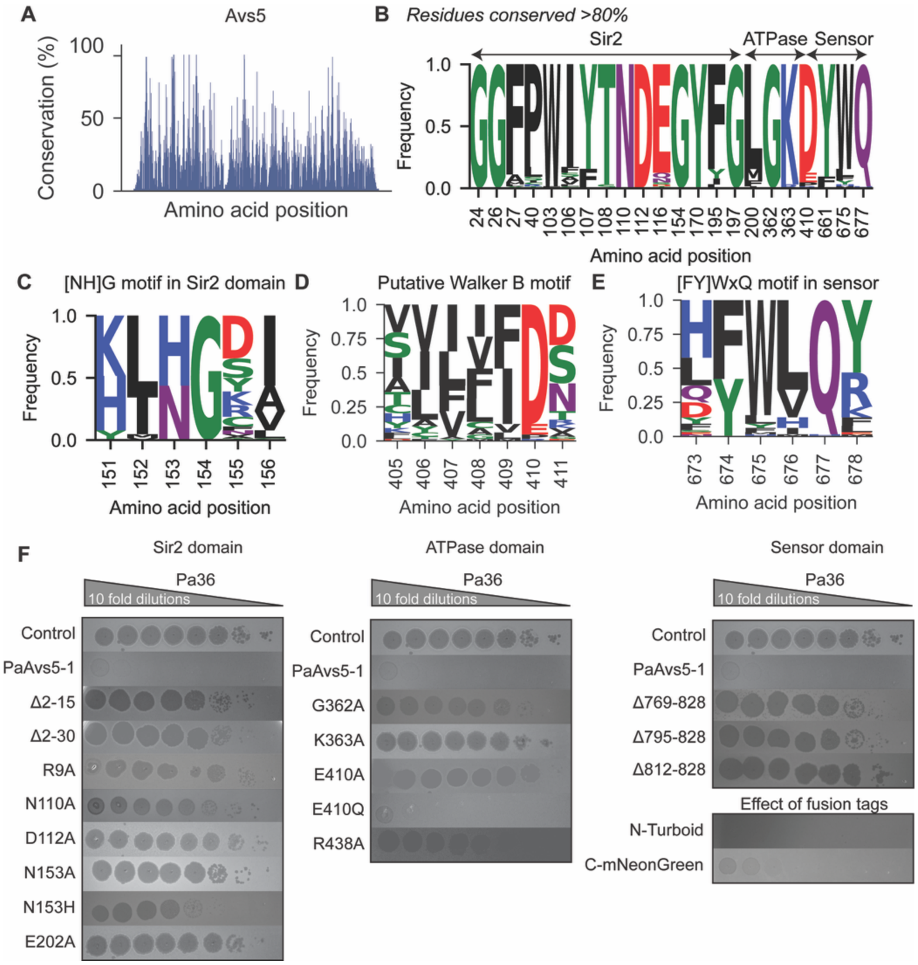


**Figure S1: Sequence Conservation and Functional Analysis of Avs5 Domains and Motifs**

(A) Plot showing percent amino acid conservation across 64 Avs5 homologs, based on multiple sequence alignment.

(B) Sequence logo highlighting residues conserved in >80% of homologs. Amino acid positions are numbered according to PaAvs5-1. Conserved regions span the Sir2, ATPase, and Sensor domains.

(C–E) Sequence logos of conserved motifs within functional domains of PaAvs5-1: (C) [NH]G motif in the Sir2 domain; (D) Putative Walker B motif in the ATPase domain; (E) [FY]WxQ motif in the sensor domain.

(F) Efficiency of plaquing (EOP) assay in *Pseudomonas aeruginosa* strain Pa36 for wild-type PaAvs5-1 and a panel of domain truncations, point mutants, and fluorescent fusion variants. Tenfold serial dilutions of phage lysate were spotted to assess antiviral activity.

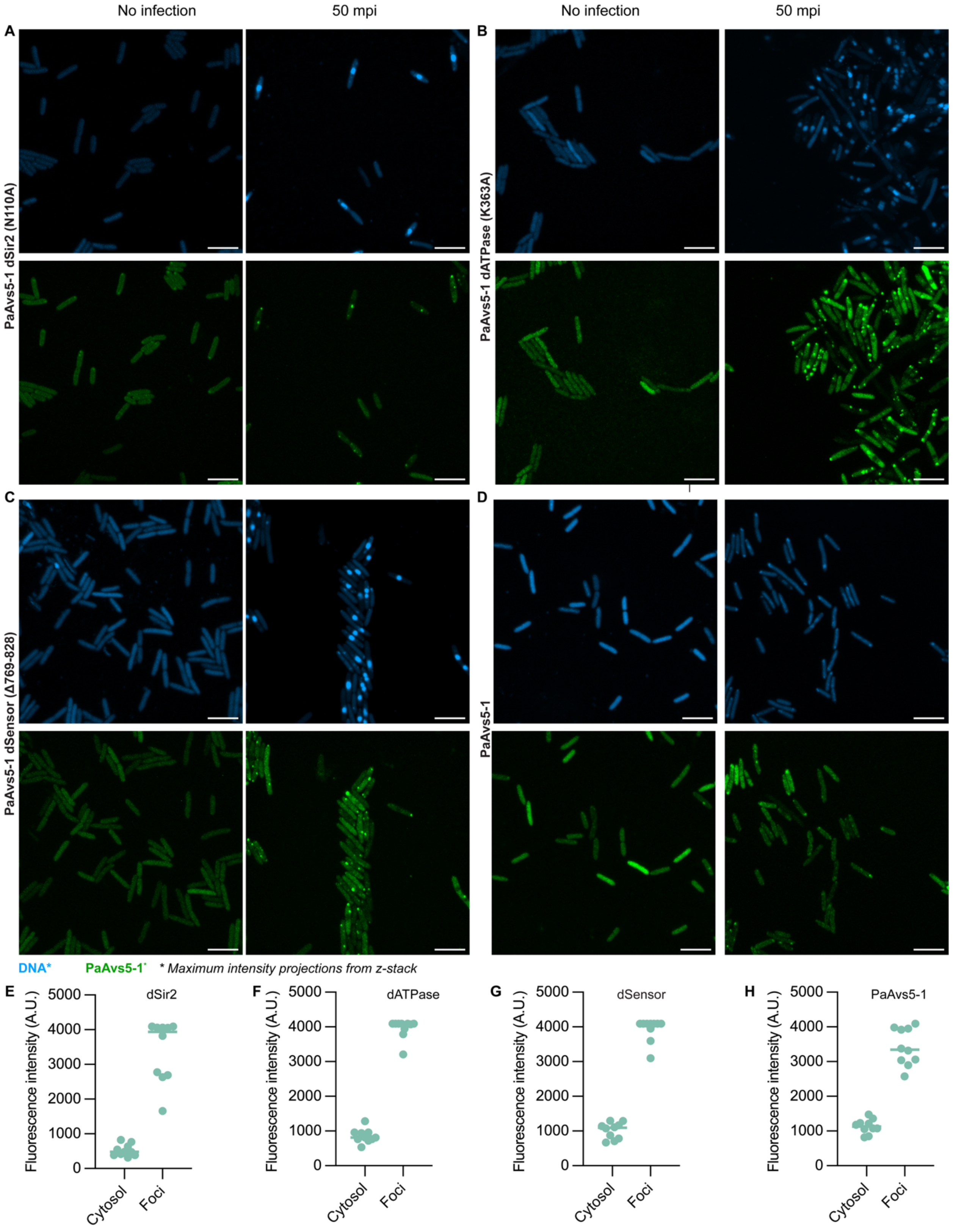


**Figure S2: Subcellular Localization of PaAvs5-1 and Domain Mutants During Phage Infection**

(A–D) Maximum intensity projection confocal fluorescence microscopy images of *P. aeruginosa* expressing wild-type or mutant PaAvs5-1 variants at indicated time points post-infection. DNA is stained in blue (DAPI), and PaAvs5-1 or its domain mutants are fused to a fluorescent tag and shown in green. (A) PaAvs5-1 dSir2 (N110A), (B) PaAvs5-1 dATPase (K363A), (C) PaAvs5-1 dSensor (Δ769–828), (D) Full-length PaAvs5-1. Scale bars represent 5 μm.

(E–H) Quantification of fluorescence intensity in the cytosol versus PaAvs5-1 foci (green channel). Ten randomly selected cells or foci were measured per sample. (E) dSir2, (F) dATPase, (G) dSensor, (H) Full-length PaAvs5-1. Maximum fluorescence intensity of 4095 AU indicates saturation.

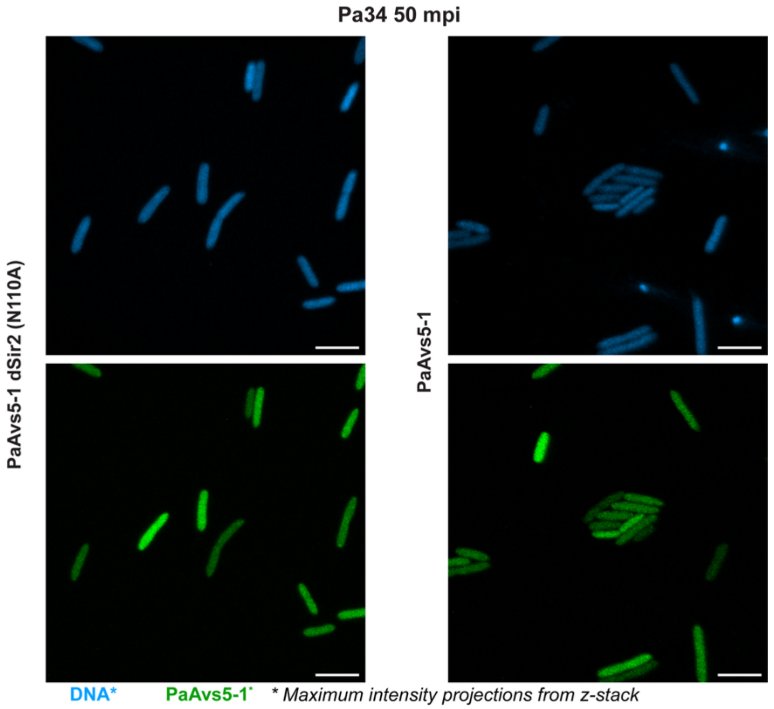


**Figure S3: Localization of PaAvs5-1 During Pa34 Infection**

Confocal fluorescence microscopy images of *P. aeruginosa* expressing PaAvs5-1 or its catalytically inactive mutant (dSir2 N110A), 50 minutes post-infection (mpi) with phage Pa34. DNA is shown in blue (DAPI), and PaAvs5-1 is shown in green. Unlike during Pa36 infection, PaAvs5-1 fails to form discrete foci during Pa34 infection, consistent with its inability to provide immunity against this phage. Scale bars: 5 μm.

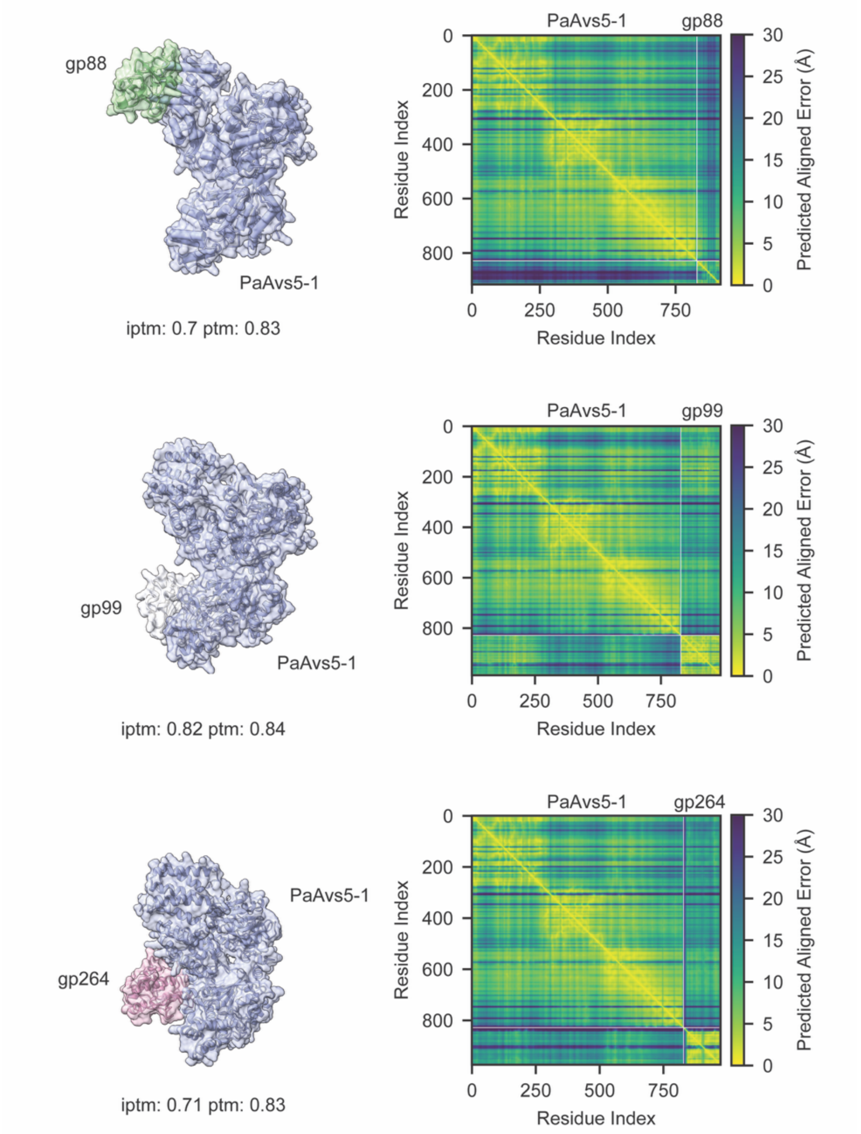


**Figure S4: AlphaFold3 Cofolding Predictions to Identify Potential Interactors of PaAvs5-1**

Structural predictions (left) and predicted aligned error (PAE) matrices (right) for PaAvs5-1 cofolded with three phage proteins from *Pseudomonas aeruginosa* phage Pa36 genome: gp88 (top), gp99 (middle), and gp264 (bottom). All 356 predicted phage proteins were tested using AlphaFold3, and these three candidates showed inter-protein TM (iptm) scores above 0.7, indicating possible stable interfaces. PAE plots reflect model confidence, with lower values (yellow) indicating higher accuracy in residue-residue alignment.

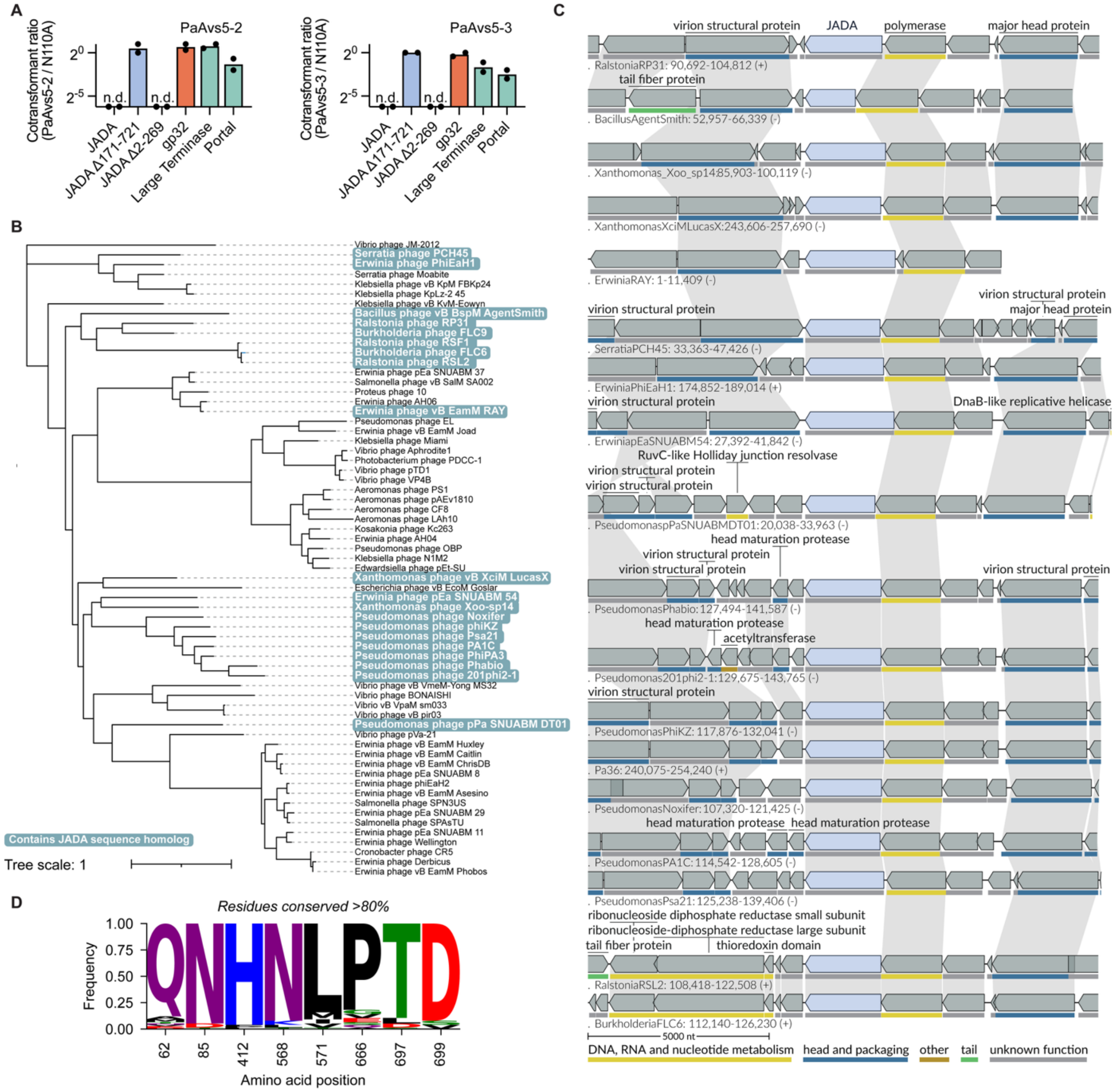


**Figure S5: Conservation and Genomic Context of JADA Homologs Across Jumbo Phages**

(A) Cotransformation assay testing activation of PaAvs5-2 and PaAvs5-3 by candidate activating genes (large terminase, portal, and JADA truncations) identified in Figure 5 of the main manuscript. Relative transformation ratios are shown for each combination, normalized to PaAvs5-1 dSir2 (N110A) control. n.d. indicates "not detectable," corresponding to zero colonies observed in transformation assays.

(B) Phylogenetic tree of nucleus-forming jumbo phages, adapted from Prichard et al. (2023) Phages encoding JADA sequence homologs are highlighted in blue.

(C) Genomic neighbourhoods of JADA homologs across representative jumbo phages. An invariant feature across all contexts is the presence of a non-virion RNA polymerase gene located immediately upstream of the JADA homolog, suggesting a conserved functional or regulatory linkage.

(D) Sequence logo showing residues conserved in >80% of JADA homologs. Residue numbering corresponds to the reference JADA protein in Pa36.

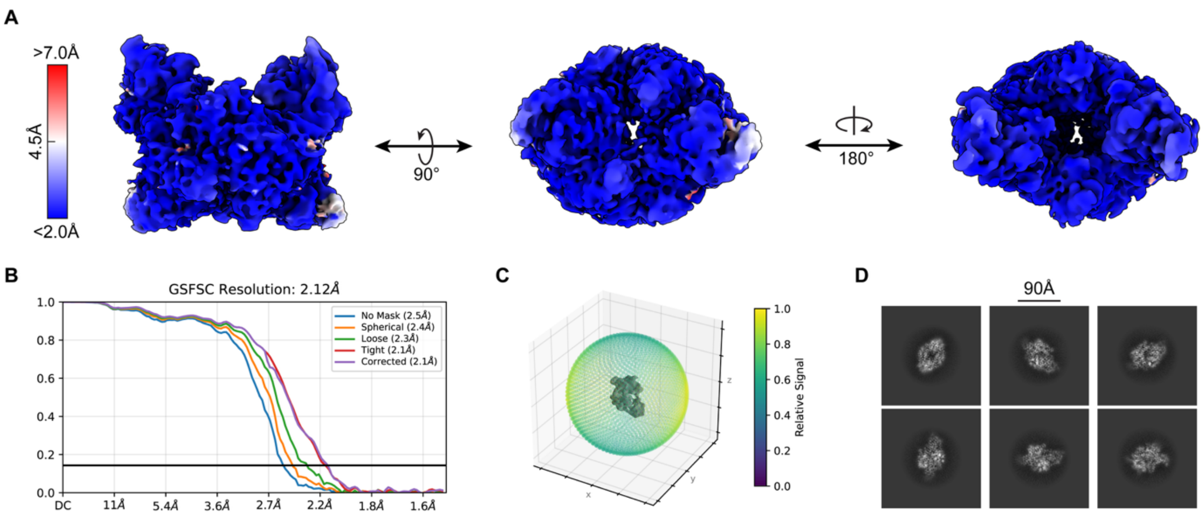


**Figure S6: Cryo-EM Map Quality and Resolution Estimation of JADA Homodimer**

(A) Local resolution estimation of the JADA cryo-EM map, displayed with color gradient from high (blue) to low (red) resolution. Three orthogonal views of the density map are shown, indicating an overall well-resolved structure.

(B) Gold-standard Fourier shell correlation (GSFSC) curves for different masking conditions, yielding an overall resolution of 2.12 Å at the 0.143 threshold.

(C) 3D directional Fourier shell correlation (dFSC) plot shows isotropic resolution across the map.

(D) Representative 2D class averages of JADA particles, highlighting structural features and particle homogeneity.

**Table S2:** Cryo-EM Data Collection, Processing, and Model Refinement Statistics for JADA/Gp316 Homodimer (PDB: 9RP3, EMDB: 54139)

| | *JADA/Gp316 homodimer (PDB: 9RP3 , EMDB: 54139)* |
|---|---|
| Voltage (kV) | 300 |
| Electron exposure (e–/Å$^2$) | 50.0 |
| Defocus range (μm) | -0.8-2.0 |
| Pixel size (Å) | 0.726 |
| Symmetry imposed | C1 |
| Initial particle images (no.) | 2,351,876 |
| Final particle images (no.) | 1,723,569 |
| Map resolution (Å) | 2.12 |
| FSC threshold | 0.143 |
| Map sharpening *B* factor (Å$^2$) | -78.0 |
| Model composition | |
| Non-hydrogen atoms | 6349 |
| Protein residues | 799 |
| Ligands | 0 |
| ADP B factors (Å$^2$) | |
| Protein | 94.1 |
| Ligand | - |
| R.m.s. deviations | |
| Bond lengths (Å) | 0.003 |
| Bond angles (°) | 0.664 |
| Validation | |
| MolProbity score | 1.46 |
| Clashscore | 8.52 |
| Poor rotamers (%) | 0.72 |
| Ramachandran plot | |
| Favored (%) | 98.74 |
| Allowed (%) | 1.26 |
| Disallowed (%) | 0.00 |

**Table S3:** List of Primers Used in This Study.

| Name | Sequence 5' to 3' | Description |
|---|---|---|
| BN5293 | P-GCTGAGCCAGCAGGGCCTGTTA | Introduce G362A mutation in PaAvs5-1 |
| BN5294 | P- AAAAACAACCACCTTAAAACAAATGGCATGGC | Introduce G362A mutation in PaAvs5-1 |
| BN5295 | P-GCTCCTGAGCCAGCAGGGCC | Introduce K362A mutation in PaAvs5-1 |
| BN5296 | P-AACAACCACCTTAAAACAAATGGCATGGCG | Introduce K362A mutation in PaAvs5-1 |
| BN4863 | GATCCTCTAGAGTCGAGCGCAT | Amplify pSTDesR FW |
| BN4864 | CCCGGGTACCGAGCTCGAAT | Amplify pSTDesR RV |
| BN4884 | P-CGCTGTTGTATATATTCTTTTCCATGAGTA | Introduce N110A mutation in PaAvs5-1 |
| BN4885 | P-ATCGATGACTCCTTTGAGAA | Introduce N110A mutation in PaAvs5-1 |
| BN4886 | P-CGCATTTATTTGTGTTCCGATAAAAAT | Introduce E202A mutation in PaAvs5-1 |
| BN4887 | P-CCGGTATTCTTCCACACTAT | Introduce E202A mutation in PaAvs5-1 |
| BN5241 | TTGAATTCATCGCTATTCATCTTTTCGGCAGACCGCAGA | Amplify turboID for N terminal fusion PaAvs5-1 |
| BN5242 | CTTTTATTTAGGAGTAAAAAATGAAAGACAATACTGTGCCTCTGAAG | Amplify turboID for N terminal fusion PaAvs5-1 |
| BN5267 | AGTCAACGCGCTGAATTAGATTACTTGTACAGCTCGTCCATGCCC | Amplify mNeonGreen for C terminal fusion PaAvs5-1 |
| BN5268 | CAACCCTAAAGCAAGAGGCAATGGTGAGCAAGGGCGAGGA | Amplify mNeonGreen for C terminal fusion PaAvs5-1 |
| BN5269 | TTTTTACTCCTAAATAAAAGCC | Amplify PaAvs5-1 plasmid for N-terminal cloning |
| BN5270 | ATGAATAGCGATGAATTCAA | Amplify PaAvs5-1 plasmid for N-terminal cloning |
| BN5271 | TGCCTCTTGCTTTAGGGTTG | Amplify PaAvs5-1 plasmid for C-terminal cloning |
| BN5272 | TCTAATTCAGCGCGTTGACTAT | Amplify PaAvs5-1 plasmid for C-terminal cloning |

| Name | Sequence 5' to 3' | Description |
|---|---|---|
| BN5450 | CGCGAATTCGAGCTCGGTACCCGGGATGCAGGATATCTTTCCGTA | Amplify JADA to introduce into pSTDesR plasmid |
| BN5451 | GGTATGCGCTCGACTCTAGAGGATCTTAACTTTCTAGATAACGGATTTG | Amplify JADA to introduce into pSTDesR plasmid |
| BN5674 | ACCATCATCACCACAGCCAGATGAACTATGCACAGTACAAT | Amplify 6xHis JADA to introduce to pACYC-Duet1 plasmid |
| BN5675 | GCCGAGCTCGAATTCGGATCTTATTTGATAAGCTTAAGTAGAAC | Amplify 6xHis JADA to introduce to pACYC-Duet1 plasmid |
| BN5676 | TGTACTTCCAATCCAATGCAATGAATAGCGATGAATTCAAAAG | Amplify Twin-Strep SUMO TEV PaAvs5-1 to introduce to p13SS plasmid |
| BN5677 | ATCCACTTCCAATGTTATTACTATGCCTCTTGCTTTAGGG | Amplify Twin-Strep SUMO TEV PaAvs5-1 to introduce to p13SS plasmid |
| BN5678 | TAATAACATTGGAAGTGGATAACGG | Amplify p13SS plasmid for cloning |
| BN5679 | TGCATTGGATTGGAAGTACA | Amplify p13SS plasmid for cloning |
| BN5680 | GATCCGAATTCGAGCTCGGC | Amplify pACYC-Duet-1 plasmid for cloning |
| BN5681 | CTGGCTGTGGTGATGATGGTG | Amplify pACYC-Duet-1 plasmid for cloning |
| BN4614 | CGCGAATTCGAGCTCGGTACCCGGGATGGACGTTATACTTTTCAATA | Amplify Pa36 large Terminase subunit to introduce in pSTDesR |
| BN4615 | GGTATGCGCTCGACTCTAGAGGATCTTAATACCAGCTATTTTTAGAATATC | Amplify Pa36 large Terminase subunit to introduce in pSTDesR |
| BN4616 | CGCGAATTCGAGCTCGGTACCCGGGATGGCTATACCAGGGGTTAA | Amplify Pa36 Portal to introduce in pSTDesR |
| BN4617 | GGTATGCGCTCGACTCTAGAGGATCTTATCCAATCTCATCTGATTCAGTT | Amplify Pa36 Portal to introduce in pSTDesR |
| BN6394 | P-CATCCCGGGTACCGAGC | Introduce JADA 2-269 deletion in pSTDesR plasmid |
| BN6395 | P-TGGGGTGCCACTGTTGAAG | Introduce JADA 2-269 deletion in pSTDesR plasmid |
| BN6206 | P-GCGGTGGTATAAATACGTTTCCAGC | Introduce N110A mutation in *E. coli* codon optimized PaAvs5-1 |

| Name | Sequence 5' to 3' | Description |
|---|---|---|
| BN6207 | P-CATCGATGATAGCTTTGAAAACACC | Introduce N110A mutation in *E. coli* codon optimized PaAvs5-1 |
| BN6179 | P-ACCACCCATACGCATCTTTTG | Introduce JADA 171-721 deletion in pSTDesR plasmid |
| BN6181 | P-TAAGATCCTCTAGAGTCGAGCG | Introduce JADA 171-721 deletion in pSTDesR plasmid |
| BN6177 | P-GTAGTTTTACAAAGTATAGATTTTCTAAAGACTCC | Introduce N153Q mutation in PaAvs5-1 |
| BN6178 | P-AGGGAGACATACAACAAACAAATAAAGG | Introduce N153Q mutation in PaAvs5-1 |
| BN6157 | P-GTATAAATATTATTTTTTTCTTACCTGG | Introduce E410Q mutation in PaAvs5-1 |
| BN6158 | P-AGCGAATTTCAATACACATA | Introduce E410Q mutation in PaAvs5-1 |
| BN6159 | TGTACTTCCAATCCAATGCAATGAACAGCGACGAATTCAA | Amplify *E. coli* codon optimized Twin-Strep SUMO TEV PaAvs5-1 to introduce to p13SS plasmid |
| BN6160 | ATCCACTTCCAATGTTATTATGCTTCTTGTTTCAGGGTCG | Amplify *E. coli* codon optimized Twin-Strep SUMO TEV PaAvs5-1 to introduce to p13SS plasmid |
| BN6142 | P-CACTTTTGAATTCATCGCTA | Introduce R9A mutation in PaAvs5-1 |
| BN6143 | P-CATTGAGAAGAAAAATGCTTGC | Introduce R9A mutation in PaAvs5-1 |
| BN6136 | AGCAGCCATCACCATCATCACCACAGCCAGATGAATAGCGATGAATTCAAAAGTAG | Amplify 6xHis PaAvs5-1 to introduce to pACYC-Duet1 plasmid |
| BN6137 | CTGCAGGCGCGCCGAGCTCGAATTCGGATCCTATGCCTCTTGCTTTAGGGT | Amplify 6xHis PaAvs5-1 to introduce to pACYC-Duet1 plasmid |
| BN6093 | P-GCTATAAATATTATTTTTTTCTTACCTGG | Introduce E410A mutation in PaAvs5-1 |
| BN6094 | P-GCGAATTTCAATACACATAG | Introduce E410A mutation in PaAvs5-1 |
| BN6095 | P-GCCTCGGCGGTTACTATAAGGAAG | Introduce R438A mutation in PaAvs5-1 |
| BN6096 | P-CGAAGGAATATGGAACTCAAGAATC | Introduce R438A mutation in PaAvs5-1 |

| Name | Sequence 5' to 3' | Description |
|---|---|---|
| BN6097 | P-CTCTATAAATATTATTTTTTTCTTACCTGG | Introduce R411A mutation in PaAvs5-1 |
| BN6098 | P-AATTTCAATACACATAGATGAGTTCATAG | Introduce R411A mutation in PaAvs5-1 |
| BN6071 | P-TGTAGTTTTACAAAGTATAGATTTTCT | Introduce N153H mutation in PaAvs5-1 |
| BN6072 | P-CGGAGACATACAACAAACAAAT | Introduce N153H mutation in PaAvs5-1 |
| BN6073 | P-CATTTTTTACTCCTAAATAAAAGCCCC | Introduce 2-15 deletion in PaAvs5-1 |
| BN6074 | P-GCCGGAGAGGTGATTCTTTT | Introduce 2-15 deletion in PaAvs5-1 |
| BN6075 | P-GCTAGTTTTACAAAGTATAGATTTTCTAAAGAC | Introduce N153A mutation in PaAvs5-1 |
| BN6076 | P-AGATACAGAAAAGCGGAGTA | Introduce 812-828 deletion in PaAvs5-1 |
| BN6077 | P-TAGTCTAATTCAGCGCGTTG | Introduce deletions in PaAvs5-1 (812-828, 795-828, 769-828) |
| BN6078 | P-ATTCTCTTTGTTCTGTAGTTTTTTAC | Introduce 795-828 deletion in PaAvs5-1 |
| BN6079 | P-TACTATAGATGTGGCGCAAG | Introduce 769-828 deletion in PaAvs5-1 |
| BN6067 | P-CATTTTTTACTCCTAAATAAAAGCC | Introduce 2-30 deletion in PaAvs5-1 |
| BN6068 | P-TCTACAAACAAATCAGGTGA | Introduce 2-30 deletion in PaAvs5-1 |
| BN6046 | CGCGAATTCGAGCTCGGTACCCGGGATGAAAGTCTTTGAAACAAG | Amplify peg.88 from Pa36 to insert into pSTDesR |
| BN6047 | GGTATGCGCTCGACTCTAGAGGATCTTAAGCAACTCTAGTGAAC | Amplify peg.88 from Pa36 to insert into pSTDesR |
| BN6048 | CGCGAATTCGAGCTCGGTACCCGGGATGACGTATAACCTAGGTATC | Amplify peg.99 from Pa36 to insert into pSTDesR |
| BN6049 | GGTATGCGCTCGACTCTAGAGGATCTTACCAACTGATTACCGTG | Amplify peg.99 from Pa36 to insert into pSTDesR |
| BN6050 | CGCGAATTCGAGCTCGGTACCCGGGATGAATAATCCAGAAACCCC | Amplify peg.264 from Pa36 to insert into pSTDesR |
| BN6051 | GGTATGCGCTCGACTCTAGAGGATCTTATCTCTGTTTGATTGATTTAATAACATC | Amplify peg.264 from Pa36 to insert into pSTDesR |

| Name | Sequence 5' to 3' | Description |
|---|---|---|
| BN6040 | GCATGCCTGCAGGTCGACTCTAGAGTCAAGTACTGCAGTTTATTC | Amplify PaAvs5-2 from MCO2344054.1 to insert into pUCP20 |
| BN6041 | GGAAACAGCTATGACCATGATTACGTAGAAGATCACATCACATTAAG | Amplify PaAvs5-2 from MCO2344054.1 to insert into pUCP20 |
| BN6042 | GCATGCCTGCAGGTCGACTCTAGAGCCCTTGAAAAATATGGTGGCTACACC | Amplify PaAvs5-3 from MCO2951615.1 to insert into pUCP20 |
| BN6043 | GGAAACAGCTATGACCATGATTACGGAGACGCCAGCGGCTAAGAATTC | Amplify PaAvs5-3 from MCO2951615.1 to insert into pUCP20 |
| BN5984 | CGCGAATTCGAGCTCGGTACCCGGGATGCCATTACCTGTAATCGC | Amplify peg.6 from Pa36 to insert into pSTDesR |
| BN5985 | GGTATGCGCTCGACTCTAGAGGATCTTAAATATTGGTAGACATGGTTTTATTC | Amplify peg.6 from Pa36 to insert into pSTDesR |
| BN5986 | CGCGAATTCGAGCTCGGTACCCGGGATGAATCTTAATCGTTATAAAGCGC | Amplify peg.26 from Pa36 to insert into pSTDesR |
| BN5987 | GGTATGCGCTCGACTCTAGAGGATCCTATCGCAGTTTACCACCTTTAAC | Amplify peg.26 from Pa36 to insert into pSTDesR |
| BN5988 | CGCGAATTCGAGCTCGGTACCCGGGATGCCTACGATTTCTCTTTCC | Amplify peg.32 from Pa36 to insert into pSTDesR |
| BN5989 | GGTATGCGCTCGACTCTAGAGGATCTTACAGTAGACGCCCCATC | Amplify peg.32 from Pa36 to insert into pSTDesR |
| BN5990 | CGCGAATTCGAGCTCGGTACCCGGGATGGCTGTTAACGAAAACGAAATC | Amplify peg.50 from Pa36 to insert into pSTDesR |
| BN5991 | GGTATGCGCTCGACTCTAGAGGATCTTAGTACCAGGTACCCGGTG | Amplify peg.50 from Pa36 to insert into pSTDesR |
| BN5992 | CGCGAATTCGAGCTCGGTACCCGGGATGACTCAATTTAACATCACCTG | Amplify peg.72 from Pa36 to insert into pSTDesR |
| BN5993 | GGTATGCGCTCGACTCTAGAGGATCTCAAAAATTAAATAGTACACCGCC | Amplify peg.72 from Pa36 to insert into pSTDesR |
| BN5994 | CGCGAATTCGAGCTCGGTACCCGGGATGTTAAAAACTATCATTAAACTAGACGG | Amplify peg.118 from Pa36 to insert into pSTDesR |
| BN5995 | GGTATGCGCTCGACTCTAGAGGATCTTAAAGAGTGCACGAACCG | Amplify peg.118 from Pa36 to insert into pSTDesR |

| Name | Sequence 5' to 3' | Description |
|---|---|---|
| BN5996 | CGCGAATTCGAGCTCGGTACCCGGGATGGCTAAGAAAAAAACAGTTC | Amplify peg.277 from Pa36 to insert into pSTDesR |
| BN5997 | GGTATGCGCTCGACTCTAGAGGATCCTATTCAGTTTCCTTGATTTCAC | Amplify peg.277 from Pa36 to insert into pSTDesR |
| BN5998 | CGCGAATTCGAGCTCGGTACCCGGGATGTTATTTAAAAAAGAAGTATGTATTGAG | Amplify peg.189 from Pa36 to insert into pSTDesR |
| BN5999 | GGTATGCGCTCGACTCTAGAGGATCCATGGCCGTTTACTAGATATTC | Amplify peg.189 from Pa36 to insert into pSTDesR |
| BN6000 | CGCGAATTCGAGCTCGGTACCCGGGATGACCAATATTGAAAAATCTGA | Amplify peg.96 from Pa36 to insert into pSTDesR |
| BN6001 | GGTATGCGCTCGACTCTAGAGGATCCTATTGTTTTGTCAATAGAAAGATC | Amplify peg.96 from Pa36 to insert into pSTDesR |
| BN6002 | CGCGAATTCGAGCTCGGTACCCGGGATGACGTATAACCTAGGTATCAC | Amplify peg.99 from Pa36 to insert into pSTDesR |
| BN6003 | GGTATGCGCTCGACTCTAGAGGATCTTACCAACTGATTACCGTGAC | Amplify peg.99 from Pa36 to insert into pSTDesR |
| BN6004 | CGCGAATTCGAGCTCGGTACCCGGGATGAAAATGAATAATAAGGTAATTG | Amplify peg.167 from Pa36 to insert into pSTDesR |
| BN6005 | GGTATGCGCTCGACTCTAGAGGATCTTATGGCAATAGTTCACTG | Amplify peg.167 from Pa36 to insert into pSTDesR |
| BN6006 | CGCGAATTCGAGCTCGGTACCCGGGATGACTGAACGTATTCATTAC | Amplify peg.220 from Pa36 to insert into pSTDesR |
| BN6007 | GGTATGCGCTCGACTCTAGAGGATCTTAACAGAATTTTTCTTGACC | Amplify peg.220 from Pa36 to insert into pSTDesR |
| BN6008 | CGCGAATTCGAGCTCGGTACCCGGGATGTGTGCAATTCCTGTAATAC | Amplify peg.224 from Pa36 to insert into pSTDesR |
| BN6009 | GGTATGCGCTCGACTCTAGAGGATCCTAATGCCCACCATATTCTC | Amplify peg.224 from Pa36 to insert into pSTDesR |
| BN6010 | CGCGAATTCGAGCTCGGTACCCGGGATGCAACCATATTCTAATTTTTCTGGTTTG | Amplify peg.261 from Pa36 to insert into pSTDesR |
| BN6011 | GGTATGCGCTCGACTCTAGAGGATCTTAACGCATGAAGCCAGGG | Amplify peg.261 from Pa36 to insert into pSTDesR |

**Table S4:** List of Plasmids Used in This Study.

| Name in this study | Code | Insert | Derived from vector | Resistance marker |
|---|---|---|---|---|
| pEmpty | pTU646 | - | pUCP20 | AmpR |
| pSTDesR-Empty | pTU679 | | pSTDesR | StrepR |
| p13SS-Empty | pTU115 | | p13SS | StrepR |
| pACYC-Empty | pTU000 | | pACYC-Duet1 | CmR |
| pAVAST_V | pTU652 | PaAvs5-1 from P. aeruginosa L0213 | pEmpty | AmpR |
| pAVAST_V-N110A | pTU724 | PaAvs5-1 N110A from P. aeruginosa L0213 | pAVAST_V | AmpR |
| pAVAST_V-D112A | pTU725 | PaAvs5-1 D112A from P. aeruginosa L0213 | pAVAST_V | AmpR |
| pAVAST_V-E202A | pTU726 | PaAvs5-1 E202A from P. aeruginosa L0213 | pAVAST_V | AmpR |
| pTerminase | pTU731 | Large terminase subunit from Pa36 | pSTDesR-Empty | StrepR |
| pPortal | pTU732 | Portal protein from Pa36 | pSTDesR-Empty | StrepR |
| p13SS-Avs5 | pTU898 | Twin-Strep SUMO Tev- PaAvs5-1 from P. aeruginosa L0213 | p13SS-Empty | StrepR |
| p13SS-Avs5-N110A | pTU899 | Twin-Strep SUMO Tev- PaAvs5-1 from P. aeruginosa L0213 N110A | p13SS-Empty | StrepR |
| p13SS-Avs5-E202A | pTU900 | Twin-Strep SUMO Tev- PaAvs5-1 from P. aeruginosa L0213 E202A | p13SS-Empty | StrepR |
| p13SS-Avs5-G362A | pTU901 | Twin-Strep SUMO Tev- PaAvs5-1 from P. aeruginosa L0213 G362A | p13SS-Empty | StrepR |
| p13SS-Avs5-K363A | pTU902 | Twin-Strep SUMO Tev- PaAvs5-1 from P. aeruginosa L0213 K363A | p13SS-Empty | StrepR |
| pJADA | pTU904 | JADA from Pa36 | pSTDesR-Empty | StrepR |
| pAVAST_V-TurboID | pTU905 | PaAvs5-1 from P. aeruginosa L0213 TurboID fusion N terminal | pAVAST_V | AmpR |
| pAVAST_V-G362A | pTU906 | PaAvs5-1 from P. aeruginosa L0213 G362A | pAVAST_V | AmpR |
| pAVAST_V-K363A | pTU907 | PaAvs5-1 from P. aeruginosa L0213 K363A | pAVAST_V | AmpR |
| pACYC_JADA | pTU1044 | 6xHis JADA from Pa36 | pACYC-Empty | CmR |
| pACYC_Avs5 | pTU1045 | 6xHis PaAvs5-1 from P. aeruginosa L0213 | pACYC-Empty | CmR |

| Name in this study | Code | Insert | Derived from vector | Resistance marker |
|---|---|---|---|---|
| pAVAST_V-delSPN | pTU1047 | Deletion 2-15 | pAVAST_V | AmpR |
| pAVAST_V-delSP | pTU1048 | Deletion 2-30 | pAVAST_V | AmpR |
| pAVAST_V-del769 | pTU1049 | Deletion 769-828 | pAVAST_V | AmpR |
| pAVAST_V-del769mNGC | pTU1050 | Deletion 769-828 mNeonGreen C terminal fusion | pAVAST_V | AmpR |
| pAVAST_V-del795 | pTU1051 | Deletion 795-828 | pAVAST_V | AmpR |
| pAVAST_V-del812 | pTU1053 | Deletion 812-828 | pAVAST_V | AmpR |
| pAVAST_V-K363AmNGC | pTU1055 | Mutation K363A mNeonGreen C terminal fusion | pAVAST_V | AmpR |
| pAVAST_V-N110AmNGC | pTU1056 | Mutation N110A mNeonGreen C terminal fusion | pAVAST_V | AmpR |
| pAVAST_V-N153H | pTU1057 | Mutation N153H | pAVAST_V | AmpR |
| pAVAST_V-N153A | pTU1058 | Mutation N153A | pAVAST_V | AmpR |
| pAVAST_V-R438A | pTU1059 | Mutation R438A | pAVAST_V | AmpR |
| pAVAST_V2 | pTU1061 | PaAvs5-2 | pEmpty | AmpR |
| pAVAST_V3 | pTU1062 | PaAvs5-3 | pEmpty | AmpR |
| pAVAST_V-TurboIDC | pTU1063 | TurboID C terminal fusion | pAVAST_V | AmpR |
| pGp88 | pTU1066 | Gp88 from Pa36 | pSTDesR-Empty | StrepR |
| pGp99 | pTU1067 | Gp99 from Pa36 | pSTDesR-Empty | StrepR |
| pGp264 | pTU1068 | Gp264 from Pa36 | pSTDesR-Empty | StrepR |
| pGp32 | pTU1073 | Gp32 from Pa36 | pSTDesR-Empty | StrepR |
| pGp50 | pTU1074 | Gp50 from Pa36 | pSTDesR-Empty | StrepR |
| pGp72 | pTU1075 | Gp72 from Pa36 | pSTDesR-Empty | StrepR |
| pGp118 | pTU1076 | Gp118 from Pa36 | pSTDesR-Empty | StrepR |
| pGp277 | pTU1077 | Gp277 from Pa36 | pSTDesR-Empty | StrepR |
| pGp189 | pTU1078 | Gp189 from Pa36 | pSTDesR-Empty | StrepR |
| pAVAST_V-R9A | pTU1096 | Mutation R9A | pAVAST_V | AmpR |

| Name in this study | Code | Insert | Derived from vector | Resistance marker |
|---|---|---|---|---|
| pAVAST_V-mNGC | pTU1098 | mNeonGreen C terminal fusion | pAVAST_V | AmpR |
| pAVAST_V-N153Q | pTU1099 | N153Q | pAVAST_V | AmpR |
| pJADA-N | pTU1100 | Deletion 171-721 | pJADA | StrepR |
| p13SS-Avs5Codon | pTU1102 | *E. coli* codon optimized Twin-Strep SUMO Tev- PaAvs5-1 from P. aeruginosa L0213 | p13SS-Empty | CmR |
| pJADA-C | | Deletion 2-269 | pJADA | StrepR |
